# Identification of MCM8IP, an interactor of MCM8-9 and RPA1 that promotes homologous recombination and DNA synthesis in response to DNA damage

**DOI:** 10.1101/751974

**Authors:** Jen-Wei Huang, Angelo Taglialatela, Ananya Acharya, Giuseppe Leuzzi, Tarun S. Nambiar, Raquel Cuella-Martin, Samuel B. Hayward, Gregory J. Brunette, Roopesh Anand, Rajesh K. Soni, Nathan L. Clark, Kara A. Bernstein, Petr Cejka, Alberto Ciccia

**Author notes:** These authors contributed equally (Angelo Taglialatela, Ananya Acharya).

## Abstract

Homologous recombination (HR) mediates the error-free repair of DNA double-strand breaks to maintain genomic stability. HR is carried out by a complex network of DNA repair factors. Here we identify C17orf53/MCM8IP, an OB-fold containing protein that binds ssDNA, as a novel DNA repair factor involved in HR. MCM8IP-deficient cells exhibit HR defects, especially in long-tract gene conversion, occurring downstream of RAD51 loading, consistent with a role for MCM8IP in HR-dependent DNA synthesis. Moreover, loss of MCM8IP confers cellular sensitivity to crosslinking agents and PARP inhibition. Importantly, we identify a direct interaction with MCM8-9, a putative helicase complex mutated in Primary Ovarian Insufficiency, that is crucial for MCM8IP’s ability to promote resistance to DNA damaging agents. In addition to its association with MCM8-9, MCM8IP also binds directly to RPA1. We show that the interactions of MCM8IP with both MCM8-9 and RPA are required to maintain replication fork progression in response to treatment with crosslinking agents. Collectively, our work identifies MCM8IP as a key regulator of DNA damage-associated DNA synthesis during DNA recombination and replication.

## MAIN TEXT

The repair of DNA double-strand breaks (DSBs) by homologous recombination (HR) is critical for genomic stability and tumor suppression^1^. HR is initiated by the nucleolytic degradation of DSBs to reveal 3’-ended single-strand DNA (ssDNA) tails, which are stabilized by the ssDNA-binding complex replication protein A (RPA)^2^. The RAD51 recombinase is then loaded onto the resected ends to form a RAD51-ssDNA nucleoprotein filament that can invade a homologous DNA duplex, resulting in the formation of a D-loop structure^3^. Within the D-loop, the invading DNA strand then primes DNA synthesis, which is catalyzed by replicative DNA polymerases in the presence of PCNA and RPA^4–6^. While the process of D-loop extension is dependent on the Pif1 helicase in yeast^7^, it remains unclear whether PIF1 and/or other DNA helicases promote DNA synthesis at D-loops in higher eukaryotes.

MCM8 and MCM9 are paralogs of the MCM2-7 replicative helicase^8^. Distinct from MCM2-7, MCM8-9 form a complex with putative DNA helicase activity that is not required for DNA replication initiation^9, 10^. Instead, the MCM8-9 complex has been implicated in HR in both mitotic and meiotic cells^11–13^. Consistently, MCM8- and MCM9-null female mice are sterile^13, 14^ and women carrying biallelic mutations in MCM8-9 exhibit Primary Ovarian Insufficiency (POI), a genetic syndrome characterized by reduced reproductive lifespan^15, 16^. Furthermore, as a consequence of their role in HR, MCM8- and MCM9-deficient cells are particularly sensitive to DNA interstrand crosslinking agents and PARP inhibition^11, 12, 17^.

MCM8-9 have been implicated in several activities both early and late in HR. MCM8-9 physically interact with and stimulate MRE11 in DSB resection^18^. MCM8-9 are also required for efficient loading of RAD51 onto DNA-damaged chromatin, potentially acting as a RAD51 mediator^11, 13^. In addition, MCM8-9 have been implicated in HR steps downstream of RAD51 loading^12, 19^, including DNA damage-induced DNA synthesis at acutely stalled replication forks, presumably the sites of one-ended DSBs, and at I-SceI-generated two-ended DSBs^19^. These observations raise the possibility that MCM8-9 can facilitate D-loop extension during recombination-associated DNA synthesis^19, 20^.

While regulation of MCM8-9 by the Fanconi anemia (FA)/BRCA pathway has been demonstrated^12, 19^, our knowledge of the physical interactors of MCM8-9 involved in HR modulation remains incomplete. In this study, we report the identification of a previously uncharacterized protein, C17orf53/MCM8IP, as a novel interactor of MCM8-9 that is recruited to sites of DNA damage in an RPA-dependent manner. Notably, MCM8IP-deficient cells exhibit HR defects downstream of RAD51 loading, especially in long-tract gene conversion, and loss of MCM8IP is associated with cellular sensitivity to cisplatin and PARP inhibition. Furthermore, MCM8IP-deficient cells exhibit slower replication fork progression and increased fork stalling in response to cisplatin. Mechanistically, we find that a direct interaction between MCM8IP and MCM8-9 is crucial for replication fork progression and cellular survival in response to crosslinking agents. Collectively, our findings support a role for the MCM8IP-MCM8-9 complex in promoting DNA damage-associated DNA synthesis.

## RESULTS

### C17orf53/MCM8IP is an RPA-associated factor

Genome stability is maintained by a complex network of factors involved in DNA damage signaling, DNA replication, recombination and repair^21^. To identify novel regulators of genome stability, we utilized the proximity-dependent biotin-identification (BioID) technology^22^. This approach employs a mutant isoform of the *E. coli* biotin ligase BirA (BirA*) that exhibits promiscuous biotin ligase activity towards all proteins in close proximity, allowing the identification of both stable and transient protein-protein interactions. For these studies, we focused our attention on RPA, a central component of the DNA damage response that interacts with a multitude of DNA replication, recombination and repair proteins^21, 23^. To identify novel RPA-associated factors, we fused BirA* to the N-terminus of RPA1, the large subunit of the RPA trimer (Figure 1A). To determine whether the BirA*-RPA1 fusion is functional, we expressed BirA*-RPA1 in U2OS cells and examined its localization in response to localized DNA damage. Following UV laser microirradiation, BirA*-RPA1 readily accumulated along the damaged tracts marked by γH2AX staining, indicating that fusion with BirA* did not impair the ability of RPA1 to localize to sites of DNA damage (Supplementary Figure 1A). Next, we expressed BirA*-RPA1 or the BirA*-alone control in HEK293T cells and performed a small-scale pulldown of biotinylated proteins using streptavidin beads in the presence or absence of hydroxyurea (HU). HU generates DSBs in S-phase after a prolonged treatment period^24^, thus being compatible with the slow labeling kinetics of BioID^22^. Relative to BirA* alone, BirA*-RPA1 was able to capture interactions with known RPA1 partners, such as RPA2 and SMARCAL1, which were further enhanced with HU treatment, indicating that the BirA* tag does not alter the interaction of RPA1 with its partners (Supplementary Figure 1B).

**Figure 1.**
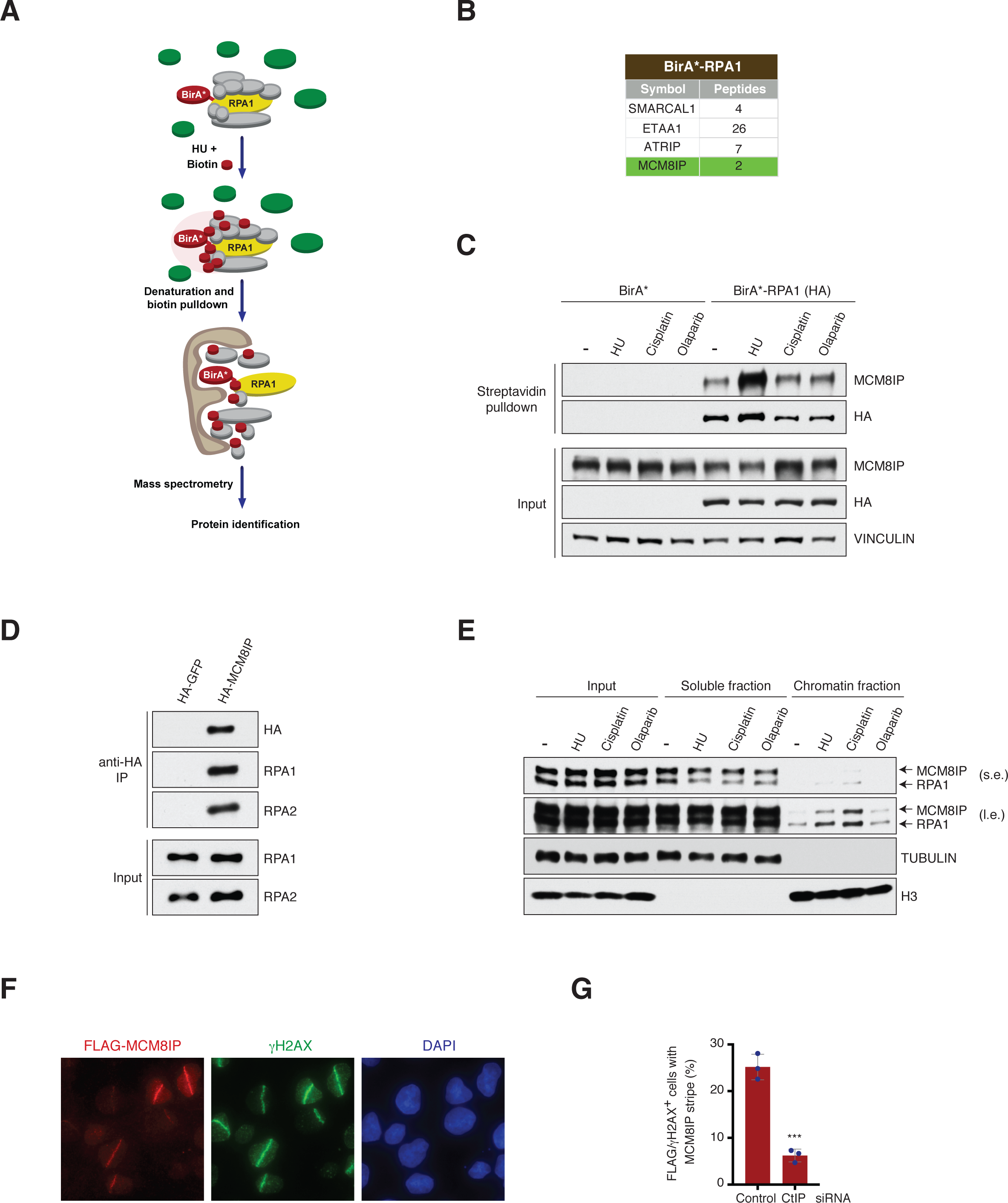
Identification of MCM8IP by proximity-dependent biotin-identification (BioID) technology. (A) Schematic of the protocol used to identify RPA1 interactors by BioID. HEK293T cells expressing BirA*- or BirA*-RPA1 were treated with HU in the presence of exogenous biotin. Biotinylated proteins were captured in denaturing conditions from cell lysates by streptavidin pulldown and subjected to mass spectrometry for protein identification. (B) List of selected proteins identified by mass spectrometry that were enriched in streptavidin pulldowns conducted from BirA*-RPA1-expressing cells relative to pulldowns performed from control BirA*-expressing cells. (C) Detection by western blot of MCM8IP in streptavidin pulldowns from HEK293T cells expressing BirA* or BirA*-RPA1. Cells were treated with HU (1 mM), cisplatin (20 µM) or olaparib (20 µM) in the presence of exogenous biotin for 24 hours prior to lysis. Vinculin is shown as a loading control. (D) Detection by western blot of RPA1 and RPA2 following immunoprecipitation of HA-GFP or HA-MCM8IP from HEK293T cells. (E) Detection by western blot of RPA1 and MCM8IP (short exposure, s.e.; long exposure, l.e.) following subcellular fractionation of HCT116 cell lysates upon treatment with HU (1 mM), cisplatin (10 µM) or olaparib (10 µM) for 24 hours. Tubulin and histone H3 are shown as loading and fractionation controls. (F) Representative images of FLAG-MCM8IP recruitment to sites of UV laser microirradiation in U2OS cells. DNA damage tracts are indicated with γH2AX staining. (G) Graphical representation of the percentage of FLAG-MCM8IP co-localizing with γH2AX following UV laser microirradiation in U2OS cells transfected with control or CtIP siRNA. The mean values ± SD of three independent experiments are presented. Statistical analysis was conducted using Student’s t-test (***p < 0.001).

Having demonstrated the functionality of BirA*-RPA1, we performed streptavidin pulldowns in HU-treated HEK293T cells expressing BirA* alone or the RPA1 fusion at scale and identified proteins by mass spectrometry. After only considering proteins represented by 2 or more peptides and at least 3-fold enriched over the control, BirA*-RPA1 identified 185 putative interactors. Several known RPA-binding proteins were identified, including ETAA1, ATRIP and SMARCAL1 (Figure 1B). Interestingly, BirA*-RPA1 captured an interaction with the previously uncharacterized C17orf53 protein, which we named MCM8IP for reasons that follow. We were able to confirm the enrichment of MCM8IP in streptavidin pulldowns from cells expressing BirA*-RPA1 by western blotting, particularly after HU treatment (Figure 1C). To validate this interaction, we conducted immunoprecipitations from cells expressing HA-tagged MCM8IP or RPA1. As shown in Figure 1D, HA-MCM8IP was able to co-immunoprecipitate RPA from HEK293T cell extracts. Importantly, HA-RPA1 reciprocally co-immunoprecipitated endogenous MCM8IP, further confirming that MCM8IP is an RPA-associated factor (Supplementary Figure 1C). These studies led to the identification of a novel interaction between RPA and MCM8IP.

### MCM8IP is recruited to chromatin after DNA damage

As the interaction between MCM8IP and BirA*-RPA1 is enhanced after treatment with DNA damaging agents (Figure 1C), we sought to determine whether MCM8IP can be recruited to sites of DNA damage. To this end, we examined bulk chromatin loading of MCM8IP by subcellular fractionation after treatment with DNA damaging agents. As shown in Figure 1E, MCM8IP was recruited to chromatin after DNA damage, especially upon treatment with HU or the crosslinking agent cisplatin. Interestingly, we noted a positive correlation between the relative amounts of chromatin-bound MCM8IP and RPA1 among the different treatments (Figure 1E). Next, we expressed FLAG-MCM8IP in U2OS cells and examined its localization in response to localized DNA damage. Following UV laser microirradiation, we observed MCM8IP accumulation onto chromatin, where it co-localized with γH2AX (Figure 1F). Interestingly, MCM8IP localization was impaired by depletion of the DSB resection-promoting factor CtIP, suggesting that it depends on the generation of 3’ ssDNA ends by DSB resection (Figure 1G and Supplementary Figure 1D-E). These findings indicate that MCM8IP accumulates at DNA damage sites containing ssDNA regions.

### MCM8IP directly interacts with RPA1

To determine whether MCM8IP and RPA1 interact directly, we partially purified GST-MCM8IP from bacteria and incubated it with lysates from bacteria expressing RPA1 or RPA2. As shown in Figure 2A, GST-MCM8IP specifically co-precipitated RPA1, whereas our positive control, GST-SMARCAL1, co-precipitated RPA2, as previously reported^25^. To identify which region of MCM8IP binds RPA1, we constructed a series of deletion mutants of GST-MCM8IP and expressed them in bacteria (Supplementary Figure 2A). Following GST pulldown of the above mutants, we identified a minimal region of MCM8IP containing the first 215 residues that was able to co-precipitate RPA1 (Supplementary Figure 2B). Several proteins that interact with RPA1 do so through acidic residue-rich motifs, which engage the basic cleft of RPA1’s N-terminal OB-fold^26–30^. Through sequence analysis, we identified two conserved stretches of acidic residues within the N-terminal third of MCM8IP (Supplementary Figure 2C). While deletion of either acidic motif from GST-MCM8IP mildly reduced the levels of co-precipitated RPA1 (Figure 2B-C), a GST-MCM8IP mutant lacking both motifs (RBM, <u>R</u>PA-<u>B</u>inding <u>M</u>utant) displayed no detectable interaction with RPA1 (Figure 2B-C). Deletion of both motifs also impaired the association of HA-MCM8IP with RPA in HEK293T cells, as determined by co-immunoprecipitation studies (Figure 2D).

**Figure 2.**
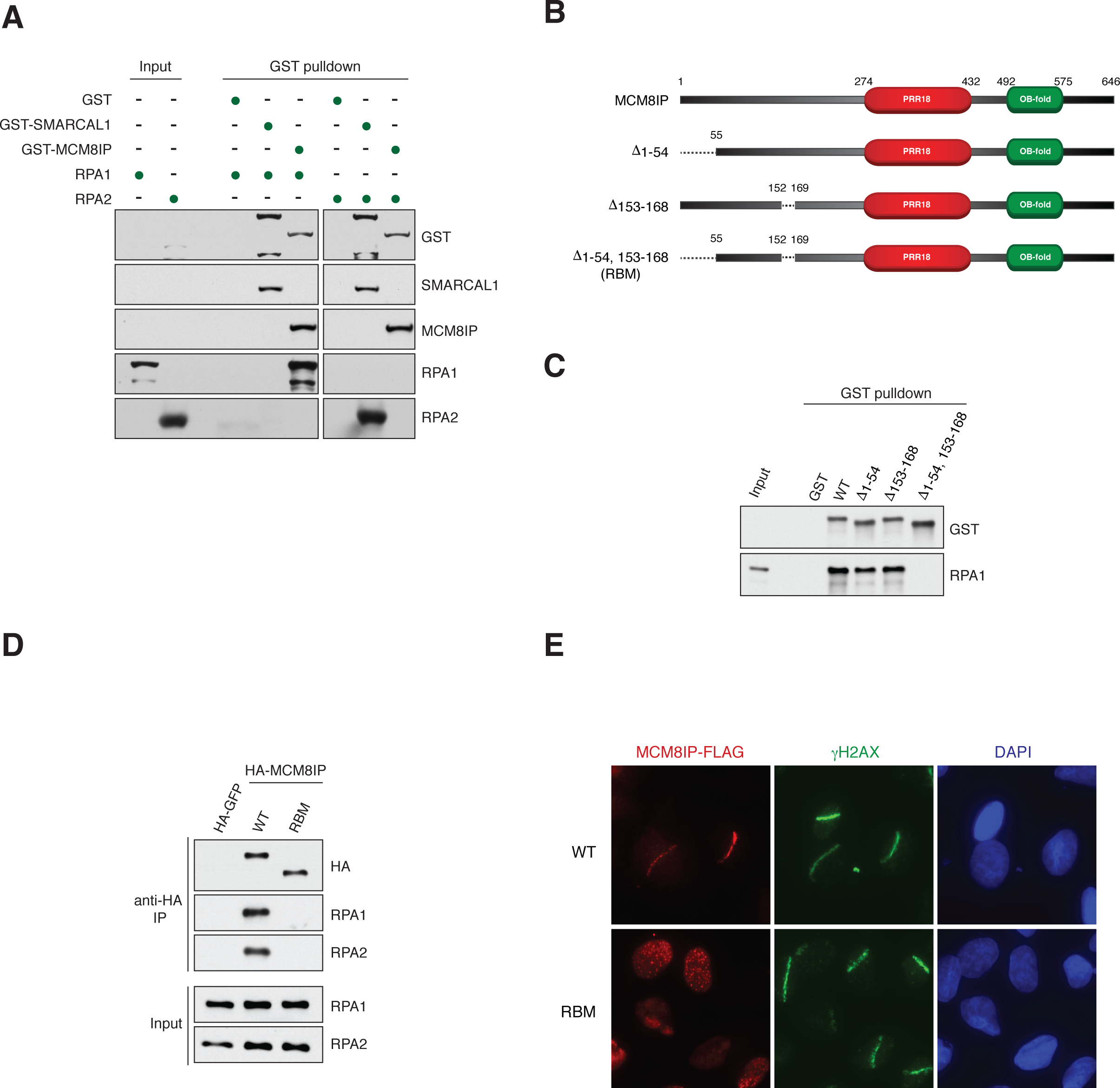
Characterization of the interaction between MCM8IP and RPA1. (A) Detection by western blot of RPA1 and RPA2 co-precipitated by bead-bound recombinant GST, GST-SMARCAL1 or GST-MCM8IP from bacteria. (B) Schematic representation of full-length MCM8IP and MCM8IP with deletions in RPA1 binding motifs. The Δ1-54,153-168 mutant was renamed RBM. Shown in red is a region with homology to Proline-rich protein 18 (PRR18). The DUF4539 domain, a predicted OB-fold, is shown in green. (C) Detection by western blot of RPA1 co-precipitated by bead-bound GST, GST-MCM8IP WT or GST fused to the MCM8IP mutants presented in (B). (D) Detection by western blot of RPA1 and RPA2 co-immunoprecipitated by HA-GFP, HA-MCM8IP WT or RBM from HEK293T cells. (E) Representative images of the recruitment of MCM8IP-FLAG WT or RBM to sites of UV laser microirradiation in U2OS cells. DNA damage tracts are indicated with γH2AX staining.

To determine whether the accumulation of MCM8IP at sites of DNA damage is dependent on its interaction with RPA, we subjected U2OS cells expressing MCM8IP-FLAG RBM to UV laser irradiation. In these experiments, MCM8IP-FLAG RBM localized to the nucleus, but failed to accumulate at sites of UV microirradiation (Figure 2E). Collectively, our findings indicate that MCM8IP contains two acidic motifs that mediate direct interaction with RPA1 and that this interaction is required for MCM8IP recruitment to DNA-damaged chromatin.

### MCM8IP interacts with MCM8-9

Although MCM8IP contains no apparent catalytic domains, it does harbor a nucleic acid-binding OB-fold motif and a central region with homology to proline-rich protein 18 (PRR18), a protein of unknown function (Figure 2B). To ascertain whether MCM8IP associates with other factors implicated in the DNA damage response, we performed streptavidin pulldowns from HU-treated cells expressing MCM8IP constructs with N- or C-terminal BirA* tags or co-immunoprecipitation from MCM8IP-HA-expressing cells. By mass spectrometric analyses, we identified a set of 22 proteins commonly found in MCM8IP BioID and co-immunoprecipitation experiments that were represented by 2 or more unique peptides and were at least 5-fold enriched in total spectral counts relative to the respective controls (Figure 3A). This set included all three subunits of RPA, as well as the RPA2-interactor, SMARCAL1. MCM8 and MCM9 were also identified among these proteins (Figure 3A), and their interaction with MCM8IP was validated by co-immunoprecipitation analyses in HEK293T cells (Figure 3B and Supplementary Figure 3A). Importantly, the MCM8IP RBM mutant co-immunoprecipitated comparable levels of MCM8-9 as WT MCM8IP, indicating that the MCM8IP-MCM8-9 interaction is independent of RPA binding (Figure 3B).

**Figure 3.**
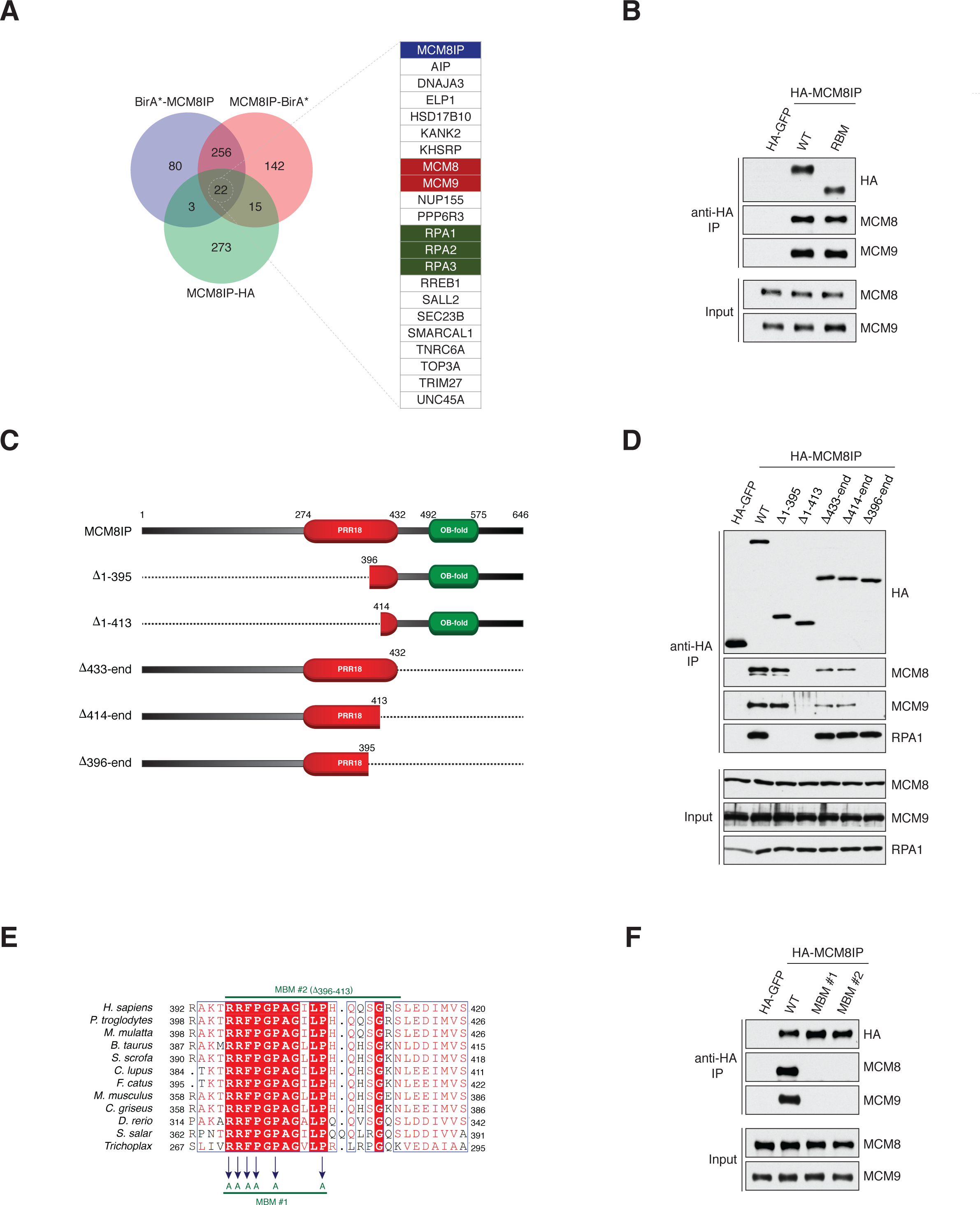
Characterization of the interaction between MCM8IP and MCM8-9. (A) Venn diagram of proteins identified by mass spectrometry that were enriched in BirA*-MCM8IP and MCM8IP-BirA* streptavidin pulldowns or MCM8IP-HA immunoprecipitates relative to their respective controls. A list of proteins commonly identified by all three experiments is presented. (B) Detection by western blot of MCM8 and MCM9 co-immunoprecipitated by HA-GFP, HA-MCM8IP WT or RBM from HEK293T cells. (C) Schematic representation of MCM8IP truncation mutants used to identify the region of interaction with MCM8-9. (D) Detection by western blot of MCM8, MCM9 and RPA1 co-immunoprepitated by HA-GFP, HA-MCM8IP WT and mutants presented in (C). (E) Alignment from various species of the minimal region of human MCM8IP required for MCM8-9 interaction. MCM8IP MBM #1 mutant carries alanine substitutions of the indicated residues. MCM8IP MBM #2 mutant carries a deletion of the indicated 18 amino acid residues. Sequence alignments were conducted using Clustal Omega and processed using ESPript. (F) Detection by western blot of MCM8 and MCM9 co-immunoprecipitated by HA-GFP, HA-MCM8IP WT, MBM #1 or MBM #2 from HEK293T cells.

Using N- and C-terminal truncation mutants of MCM8IP (Figure 3C), we identified a region spanning residues 396-413 that was sufficient for MCM8-9 interaction (Figure 3D). This region of MCM8IP contains several residues that are conserved among metazoans (Figure 3E). Substitution of six of the conserved residues with alanines (<u>M</u>CM8-9 <u>B</u>inding <u>M</u>utant #1, MBM #1) or deletion of the entire region (MBM #2) led to abrogation of the interaction of MCM8IP with MCM8-9 (Figure 3E-F). Consistently, we observed that recombinant MCM8IP interacts directly with MCM8, either alone or in complex with MCM9, and this interaction is disrupted in the MBM #1 mutant (Supplementary Figure 3B-C). In further support of an evolutionarily conserved interaction between MCM8IP and MCM8-9, we find that the *MCM8IP*, *MCM8* and *MCM9* genes significantly co-occur among multicellular eukaryotes (p-values: 3.48 x 10^-5^ for MCM8-MCM9; 1.69 x 10^-4^ for MCM8-MCM8IP; 1.03 x 10^-3^ for MCM9-MCM8IP), suggesting that MCM8IP is dependent on MCM8-9 to perform its function (Supplementary Figure 3D).

To determine whether the recruitment of MCM8IP to sites of DNA damage is dependent on its association with MCM8-9, U2OS cells expressing MCM8IP-FLAG MBM #2 were subjected to UV laser microirradiation. As shown in Supplementary Figure 3E-F, MCM8IP-FLAG MBM #2 was as efficiently recruited to sites of UV laser microirradiation as WT MCM8IP, indicating that the interaction with MCM8-9 is not required for MCM8IP recruitment to sites of DNA damage.

### The MCM8IP-MCM8-9 complex interacts with ssDNA

Our previous observations indicate that MCM8IP associates with ssDNA-containing regions in mammalian cells (Figure 1G and Supplementary Figure 1D-E). To determine whether MCM8IP directly binds ssDNA, we purified FLAG-tagged MCM8IP (Supplementary Figure 4A) from insect cells and incubated it with ^32^P-labeled ssDNA. As shown in Figure 4A-B, MCM8IP associates with ssDNA at higher concentrations (≥32 nM). Likewise, recombinant MCM8-9 complex (20 nM) purified from insect cells (Supplementary Figure 4B) also exhibits ssDNA-binding activity (Figure 4C, lane b, left panel). To determine whether the MCM8IP-MCM8-9 complex also binds ssDNA, we incubated ssDNA with MCM8-9 (20 nM) and either the WT or MBM #1 mutant form of MCM8IP (20 nM). As shown in Figure 4C, addition of WT, but not MBM #1, MCM8IP led to the formation of a protein-ssDNA complex with slower electrophoretic migration (compare lanes d with b and f, both panels), indicating that the MCM8IP-MCM8-9 complex possesses ssDNA-binding activity.

**Figure 4.**
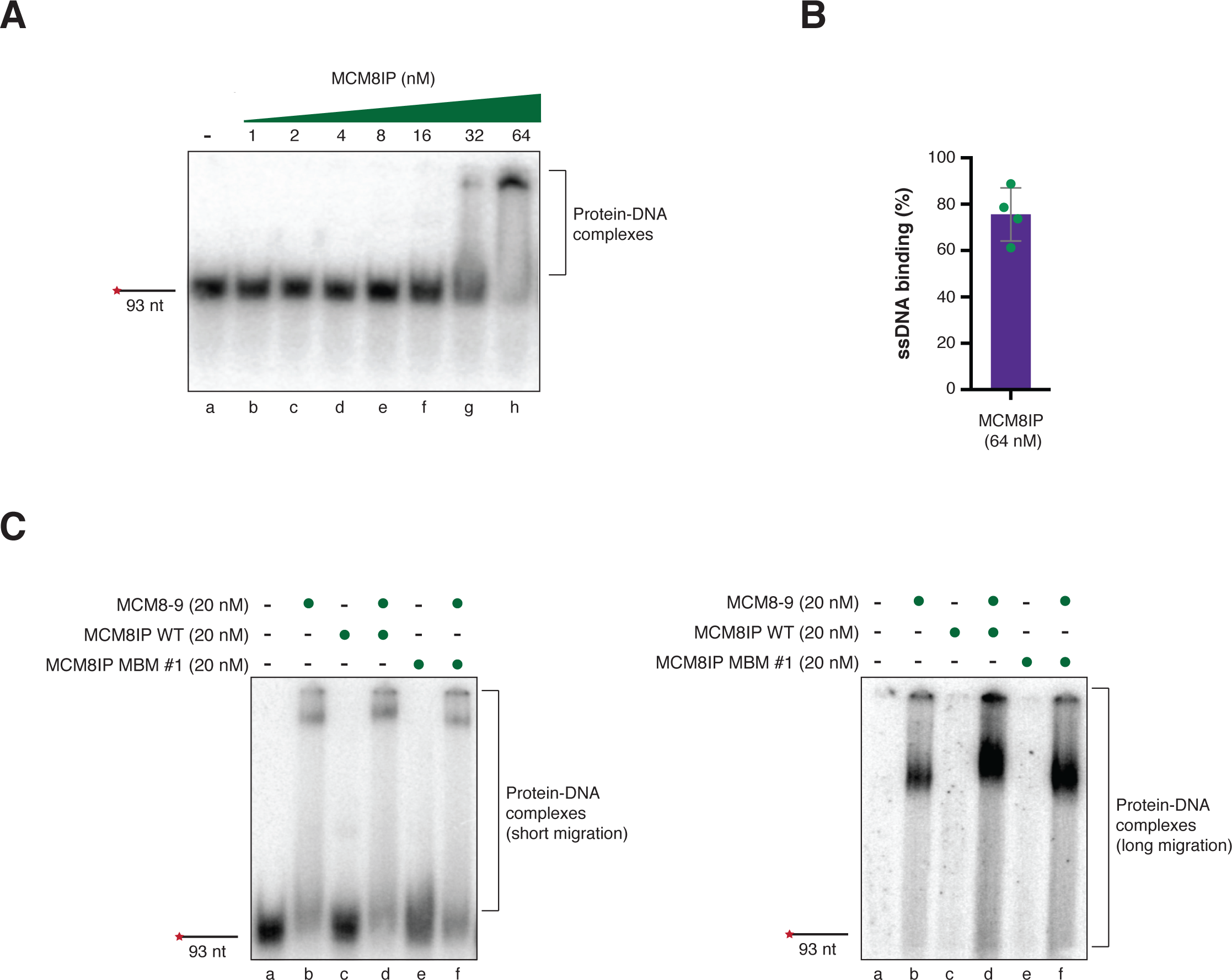
ssDNA binding properties of MCM8IP and MCM8-9. (A) Representative autoradiograph of an electrophoretic mobility shift assay with ^32^P-labeled single-stranded 93-mer incubated with increasing amounts of recombinant MCM8IP. (B) Graphical representation of the percentage of single-stranded 93-mer substrate bound by recombinant MCM8IP (64 nM) in electrophoretic mobility shift assays conducted as shown in (A). The mean ± SD of four independent experiments is presented. (C) Representative autoradiograph of an electrophoretic mobility shift assay with single-stranded 93-mer as in (A) incubated with recombinant MCM8-9 (20 nM), MCM8IP WT (20 nM) or MBM #1 (20 nM), either alone or in combination, as indicated (left panel). Longer migration of the electrophoretic mobility shift assay is shown (right panel).

### MCM8IP regulates homologous recombination

Since *MCM8IP* co-evolved significantly with genes involved in the FA and HR repair pathways (Supplementary Figure 5A), we sought to determine whether MCM8IP regulates HR. To this end, we targeted MCM8IP with three different sgRNAs using CRISPR-Cas9 in U2OS cells carrying the HR reporter construct DR-GFP (Figure 5A and Supplementary Figure 5B)^31^. Following induction of an I-SceI-mediated DSB in DR-GFP, we observed reductions in the efficiency of gene conversion, as determined by measuring the percentage of GFP-positive cells (Figure 5B). Impaired HR activity was also observed when MCM8IP was disrupted with 3 different sgRNAs in HEK293T cells carrying a BFP-based reporter of Cas9-mediated HR (Supplementary Figure 5C-E)^32^, which can be converted into GFP upon Cas9-mediated cleavage and recombination with a dsDNA donor template carrying a single nucleotide substitution (c.197C>T, p.His66Tyr)^33, 34^.

**Figure 5.**
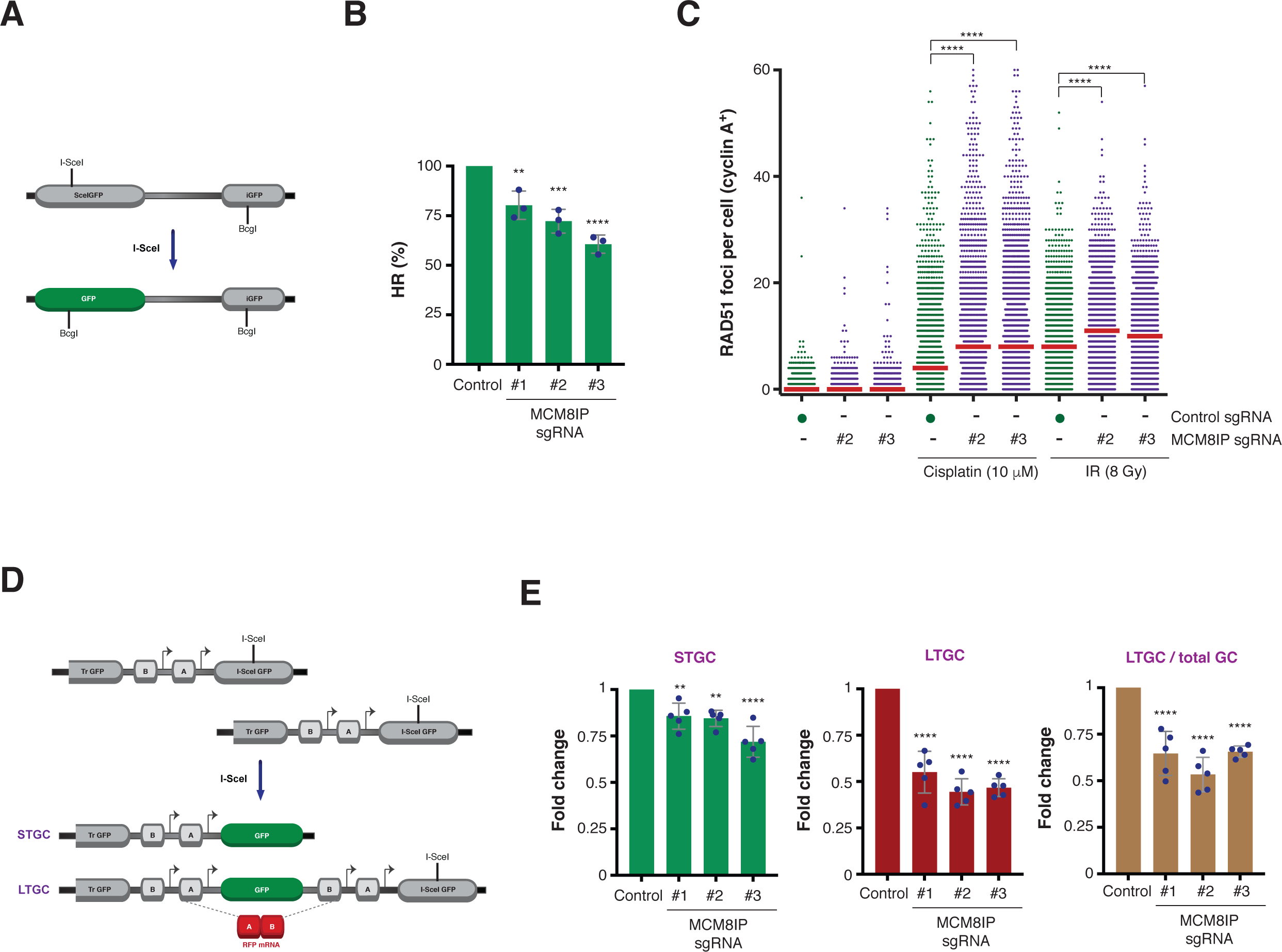
Homologous recombination frequency in MCM8IP-deficient cells. (A) Schematic representation of the DR-GFP reporter of gene conversion, as previously described^83^. Repair of an I-SceI-induced DSB in *SceGFP* by using *iGFP* as a homologous donor results in GFP expression. (B) Graphical representation of the percentage of HR events in U2OS DR-GFP cells expressing the indicated MCM8IP sgRNAs relative to the non-targeting control sgRNA. The mean ± SD of three independent experiments is presented. Statistical analysis relative to the control was conducted using one-way ANOVA (**p < 0.01, ***p < 0.001, ****p < 0.0001). (C) Dot plot of the number of cisplatin- or ionizing radiation-induced RAD51 foci in cyclin A-positive HCT116 cells expressing the indicated MCM8IP sgRNAs or a non-targeting control sgRNA. Cells were fixed after 24 hours of cisplatin treatment (10 µM) or after 6 hours from the IR treatment (8 Gy). RAD51 foci and cyclin A were stained for immunofluorescence, imaged by high-throughput microscopy and subjected to software-based quantitation. The median values are indicated by red lines. Data are representative of two independent experiments. Statistical analysis was conducted using a Mann Whitney test (****p < 0.0001). (D) Schematic representation of the SCR/RFP reporter of gene conversion, as previously described^35^. Blocks “A” and “B” are 5’ and 3’ artificial *RFP* exons. Arrows are promoters. Repair of an I-SceI induced DSB in *I-SceI GFP* by using *Tr GFP* as a homologous donor results in GFP expression. Short-tract gene conversion (STGC) results in GFP expression alone while long-tract gene conversion (LTGC) results in the additional expression of RFP. (E) Graphical representation of the fold change in STGC (left panel), LTGC (middle panel) and LTGC as a fraction of total gene conversion events (right panel) in U2OS 35S cells expressing the indicated MCM8IP sgRNAs relative to the non-targeting control sgRNA. The mean ± SD of five independent experiments is presented. Statistical analysis was conducted as in (B).

To determine the underlying defect in HR, we first examined the efficiency of DNA damage-induced RAD51 foci formation upon MCM8IP loss. In HCT116 cells, MCM8IP-depletion significantly increased RAD51 foci formation in response to either cisplatin or ionizing radiation, indicative of an HR defect downstream of RAD51-mediated strand invasion (Figures 5C and 6B). To investigate the possible role of MCM8IP in HR steps following RAD51-mediated strand invasion, we evaluated the effect of MCM8IP loss on long-tract gene conversion (LTGC), which represents a measure of recombination-associated DNA synthesis^19^. To this end, we targeted MCM8IP in 35S cells, a line of U2OS cells that carries the SCR/RFP reporter^35^. Following I-SceI-induced DSB formation in an inactive GFP gene, LTGC restores the GFP open reading frame and duplicates an RFP cassette, leading to RFP expression (Figure 5D). LTGC can therefore be evaluated by the percentage of GFP/RFP double-positive cells. Consistent with the finding from U2OS DR-GFP cells, we observed a general decrease in gene conversion in U2OS 35S cells upon MCM8IP loss (Figure 5E and Supplementary Figure 5F). Notably, however, MCM8IP loss induced a reduction of LTGC events among total GC events, indicating that MCM8IP is primarily needed for GC events that require extensive DNA synthesis. Collectively, these findings suggest that MCM8IP can promote DSB repair through regulation of HR-associated DNA synthesis.

**Figure 6.**
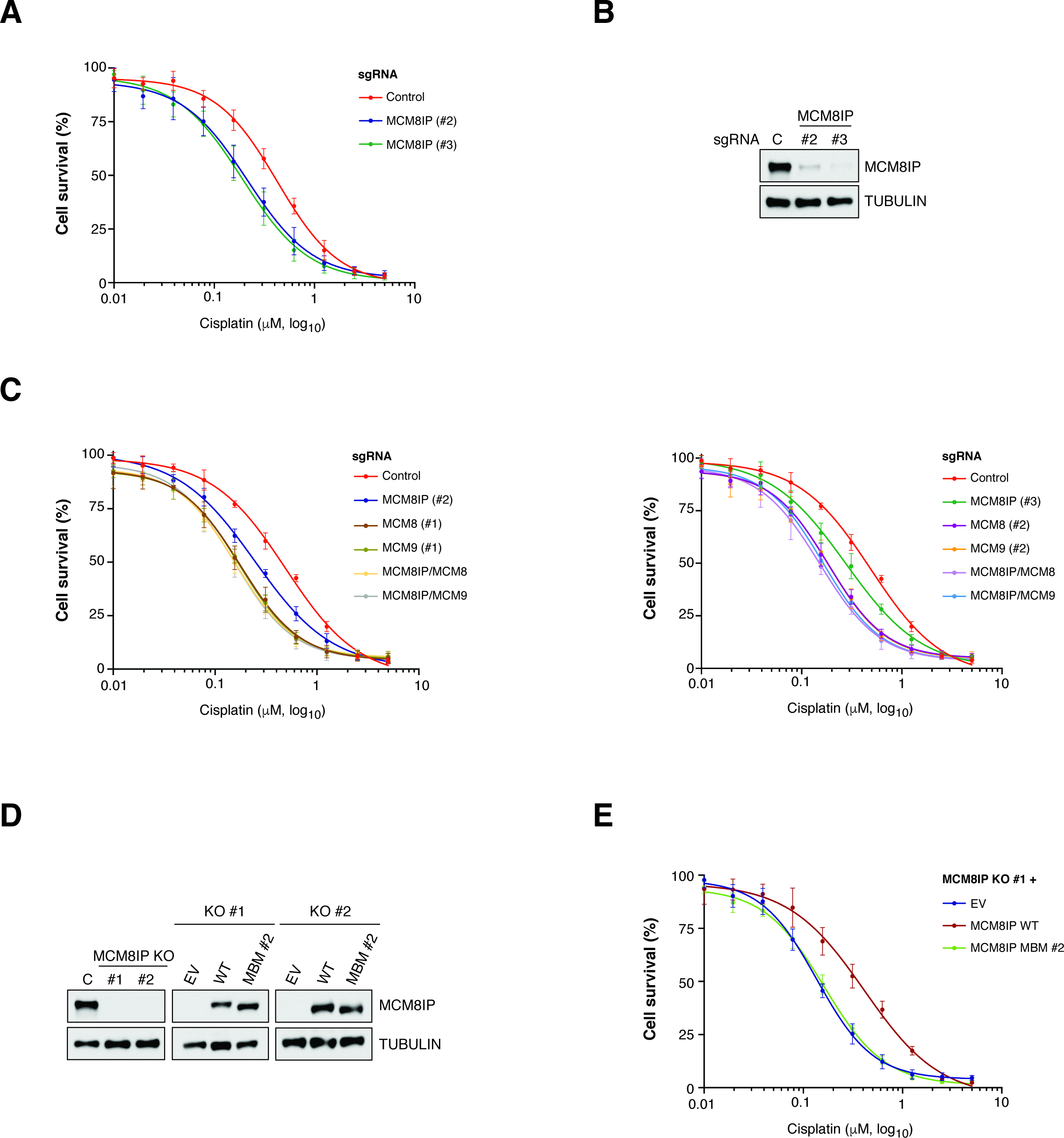
Survival analysis in MCM8IP-deficient cells treated with cisplatin. (A) Survival analysis in HCT116 cells expressing the indicated MCM8IP sgRNAs or a non-targeting control sgRNA upon treatment with cisplatin. Cell survival is expressed as a percentage of an untreated control. The mean ± SD of three independent experiments is presented. (B) Detection by western blot of MCM8IP in HCT116 cells expressing the indicated MCM8IP sgRNAs or the non-targeting control sgRNA utilized in the assay shown in (A). Tubulin is shown as a loading control. (C) Survival analysis of cisplatin-treated HCT116 control cells or cells expressing either MCM8IP sgRNA #2, MCM8 sgRNA #1, MCM9 sgRNA #1 (left panel) or MCM8IP sgRNA #3, MCM8 sgRNA #2, MCM9 sgRNA #2 (right panel), alone or in combination. Cell survival is expressed as a percentage of an untreated control. The mean ± SD of three or more independent experiments is presented. (D) Detection by western blot of MCM8IP in HCT116 MCM8IP KO clone #1, clone #2 or a non-targeting control (left panel), and in MCM8IP KO #1 (middle panel) or KO #2 (right panel) cells reconstituted with MCM8IP WT, MBM #2 or an empty vector (EV) control. Tubulin is shown as a loading control. (E) Survival analysis in HCT116 MCM8IP KO #1 cells reconstituted with MCM8IP WT, MBM #2 or an EV in response to cisplatin. Cell survival is expressed as a percentage of an untreated control. The mean ± SD of four independent experiments is presented.

### MCM8IP mediates cellular resistance to DNA damaging agents

A hallmark of HR-deficient cells is hypersensitivity to DNA interstrand-crosslinking agents^36^. We therefore sought to determine whether the HR defects of MCM8IP-deficient cells are associated with a vulnerability to cisplatin-induced cell death. To this end, we subjected HCT116 cells expressing MCM8IP sgRNAs (#2 and #3) to varying doses of cisplatin (Figure 6A-B). As shown in Figure 6A, we observed a notable reduction in cell survival of MCM8IP-deficient cells following cisplatin treatment relative to the control. These results indicate that MCM8IP-deficient cells exhibit enhanced sensitivity to treatment with crosslinking agents.

MCM8-9 deficiency is characterized by sensitivity to crosslinking agents in various species^11–13^, raising the possibility that MCM8IP cooperates with MCM8-9 to promote chemoresistance. We therefore examined whether MCM8IP and MCM8-9 genetically interact. To this end, we targeted MCM8 or MCM9 with sgRNAs in HCT116 cells expressing control sgRNA or MCM8IP sgRNAs (#2 and #3) (Supplementary Figure 6A). As expected, loss of either MCM8 or MCM9 significantly sensitized control HCT116 cells to cisplatin (Figure 6C). Additionally, we found that MCM8 or MCM9 loss also sensitized cells to PARP inhibition by olaparib (Supplementary Figure 6B), as previously observed^17^. Importantly, targeting of MCM8 or MCM9 did not further sensitize cells expressing either MCM8IP sgRNAs to cisplatin or olaparib (Figure 6C and Supplementary Figure 6B). These results suggest an epistatic relationship between MCM8IP and MCM8-9 with regard to cellular resistance to cisplatin or olaparib.

Next, we sought to determine whether the physical interaction between MCM8IP and MCM8-9 is required for chemoresistance. We derived clones from HCT116 cells expressing sgRNA #2 (KO #1) and #3 (KO #2) and confirmed the sensitivity of each clone to cisplatin and olaparib (Supplementary Figure 6C). Interestingly, expression of WT, but not MBM #2 mutant, MCM8IP cDNA restored chemoresistance in both knockout clones (Figure 6D-E and Supplementary Figure 6D-E). These results indicate that interaction with MCM8-9 is critical for MCM8IP-dependent cellular resistance to DNA damaging agents.

### MCM8IP is required for DNA synthesis in response to DNA damage

DNA interstrand crosslinks act as physical blocks to replication fork progression^37, 38^. Given the hypersensitivity of MCM8IP-deficient cells to crosslinking agents, we examined whether MCM8IP deficiency affects replication fork progression upon cisplatin treatment using the DNA fiber assay (Figure 7A). In untreated conditions, HCT116 cells expressing either control or MCM8IP sgRNAs exhibited similar rates of DNA synthesis, as indicated by the comparable lengths of IdU-labeled tracts (Figure 7A-B). While the addition of cisplatin only mildly reduced fork progression in HCT116 control cells, cells expressing either MCM8IP sgRNA exhibited significantly shorter tracts of DNA synthesis (Figure 7A-B). Consistently, we also observed increased cisplatin-induced fork stalling in MCM8IP-deficient cells (Supplementary Figure 7A). Collectively, these results indicate that MCM8IP is required for efficient DNA synthesis in the presence of cisplatin.

**Figure 7.**
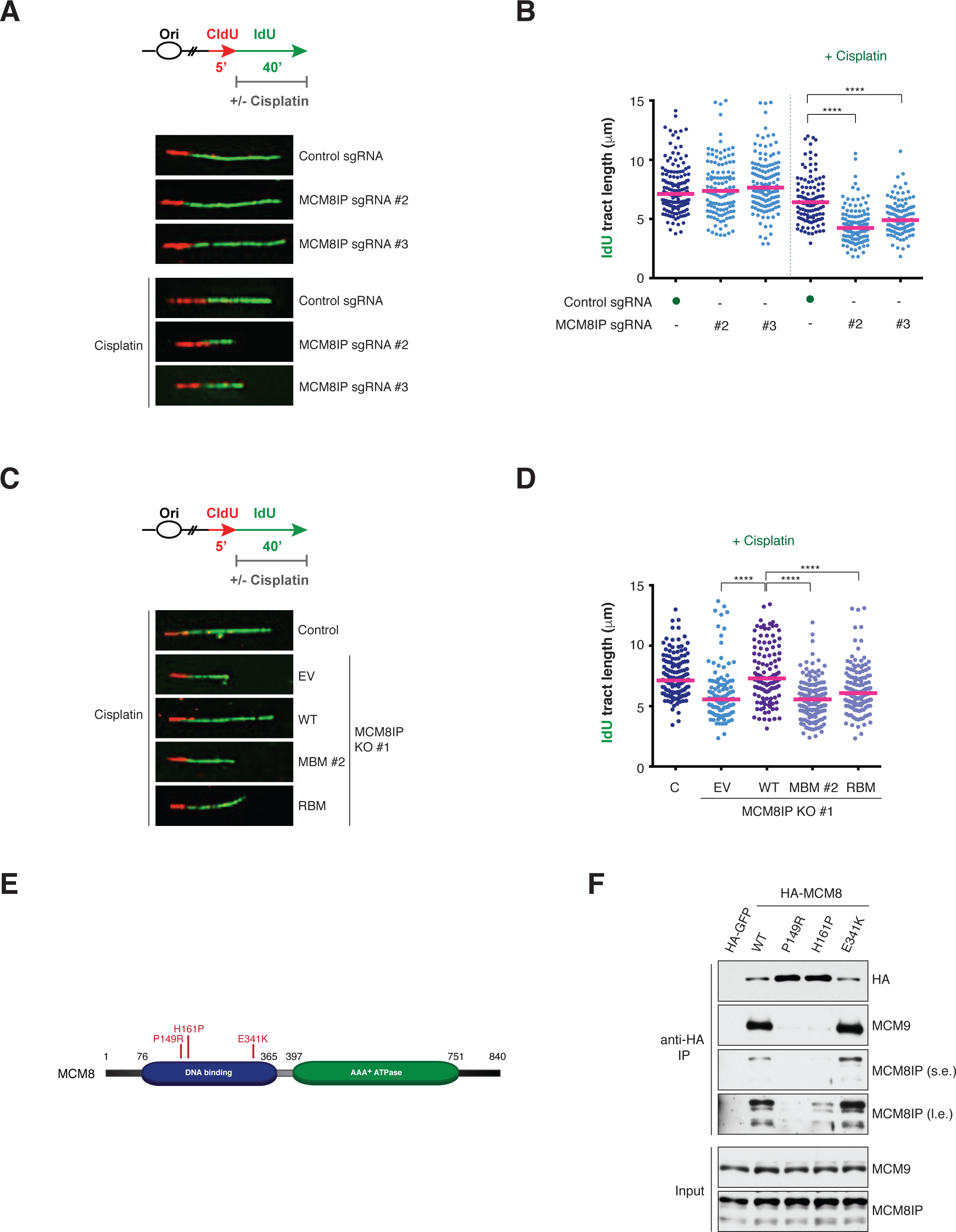
Analysis of replication fork progression in MCM8IP-deficient cells in response to cisplatin treatment. (A) Schematic representation of the CldU/IdU pulse-labeling assay (top panel). IdU labeling was performed in the absence or presence of cisplatin (30 µM). Representative images of fibers analyzed in untreated or cisplatin-treated HCT116 cells expressing the indicated MCM8IP sgRNAs or a non-targeting control sgRNA (bottom panel). (B) Dot plot of IdU tract length for individual replication forks in untreated or cisplatin-treated HCT116 cells expressing the indicated MCM8IP sgRNAs or a non-targeting control sgRNA. Experiments were conducted as shown in (A). The median values are indicated by red lines. Statistical analysis was conducted using a Mann Whitney test (****p < 0.0001). Data are representative of two independent experiments. (C) Schematic representation of the CldU/IdU pulse-labeling assay as in (A) (top panel). Representative images of fibers analyzed in cisplatin-treated HCT116 non-targeting control cells or MCM8IP KO #1 cells reconstituted with MCM8IP WT, MBM #2, RBM or an empty vector (EV) control (bottom panel). (D) Dot plot of IdU tract length for individual replication forks in cisplatin-treated HCT116 non-targeting control cells or MCM8IP KO #1 cells reconstituted with MCM8IP WT, MBM #2, RBM or EV. Experiments were conducted as shown in (C). Data are shown and analyzed as in (B) and are representative of two independent experiments. (E) Schematic representation of MCM8 with POI-associated mutations indicated. The DNA binding domain is indicated in blue and the ATPase domain is indicated in green. (F) Detection by western blot of MCM8IP (short exposure, s.e.; long exposure, l.e.) and MCM9 co-immunoprecipitated by HA-GFP, HA-MCM8 WT or carrying the POI-associated mutations indicated in (E).

We next examined which MCM8IP interactions are required to maintain fork progression on cisplatin-damaged DNA by monitoring fork progression in our MCM8IP KO #1 clone expressing either the WT, MBM #2 mutant, or RBM mutant form of MCM8IP (Figure 7C-D). Consistent with our earlier results (Figure 7B), replication fork progression was comparable between MCM8IP KO #1 and control sgRNA-expressing cells in unperturbed conditions (Supplementary Figure 7B). Importantly, the addition of cisplatin strongly impeded fork progression in MCM8IP KO #1 cells relative to the control, and this effect was largely rescued by the expression of WT MCM8IP cDNA (Figure 7D). Intriguingly, neither MBM #2 nor the RBM mutant was able to restore proper fork progression upon cisplatin treatment (Figure 7D and Supplementary Figure 7C). These findings indicate that interactions with both RPA and MCM8-9 are required for MCM8IP to maintain proper replication fork progression in the presence of cisplatin-damaged DNA.

### POI patient mutations disrupt the MCM8IP-MCM8 interaction

Biallelic mutations in *MCM8* predispose women to Primary Ovarian Insufficiency (POI)^16^. This phenotype is consistent with the sterility of female MCM8-null mice, which exhibit defects in gametogenesis due to impaired HR during meiosis^13^. Having demonstrated the functional cooperation between MCM8IP and MCM8-9, we examined whether POI-associated MCM8 mutations (i.e., P149R, H161P, E341K, Figure 7E)^39–42^ could disrupt the interaction of MCM8 with MCM8IP. To this end, we expressed either WT or mutant HA-MCM8 in HEK293T cells and subjected the cell lysates to anti-HA immunoprecipitation. As shown in Figure 7F, MCM8 carrying P149R or H161P mutations co-immunoprecipitated significantly less MCM8IP and MCM9 than the WT control or MCM8 E341K. These results indicate that disruption of the MCM8IP-MCM8-MCM9 complex may underlie the etiology of POI in patients carrying certain MCM8 mutations.

## DISCUSSION

Our study implicates the previously uncharacterized C17orf53 protein and its interaction with MCM8-9 in the regulation of HR. Accordingly, we refer to C17orf53 as MCM8IP (<u>MCM8</u>-9-Interacting <u>P</u>rotein). We find that the HR defect caused by MCM8IP deficiency is consistent with an impairment in recombination-associated DNA synthesis downstream of RAD51 loading (Figure 5C-E), as in the case of MCM8-9 deficient cells^19^. We also demonstrate that the interaction between MCM8IP and MCM8-9 is important to mediate cellular resistance to DNA damaging agents. In addition to interacting with MCM8-9, we show that MCM8IP directly associates with RPA. Both the MCM8IP-RPA and MCM8IP-MCM8-9 interactions are important for maintaining replication fork progression in response to DNA damage induced by crosslinking agents. Collectively, these findings highlight the importance of MCM8IP in promoting DNA damage-associated DNA synthesis at replication and recombination intermediates.

### MCM8IP and the regulation of HR

Previous studies of MCM8-9 indicate a multifaceted role both early and late in HR^18, 19^, with the multiple functions potentially regulated through distinct and/or mutually exclusive MCM8-9-containing complexes. For example, MCM8-9 may regulate DSB resection early in HR upon association with the MRE11-RAD50-NBS1 (MRN) complex^18^. Here we propose that the MCM8IP-MCM8-9 complex promotes recombination-associated DNA synthesis late in HR. Thus, the composition of different MCM8-9-containing complexes and how they are regulated adds an intriguing layer of regulation to HR. Future proteomic analyses of MCM8-9 binding partners, particularly in the context of DNA damage, will provide a more comprehensive understanding of the functions of MCM8-9 in HR and genome stability.

As an alternative replicative helicase, MCM8-9 has been proposed to drive D-loop extension and migration during gene conversion and break-induced replication (BIR)^19^. Our study indicates that interaction with MCM8IP is required for MCM8-9 function. The exact nature of the regulation of MCM8-9 by MCM8IP remains to be fully elucidated. One potential function of MCM8IP could be in determining the substrate specificity of the MCM8IP-MCM8-9 complex. Interaction with MCM8-9 may, for example, confer or enhance the affinity of MCM8-9 for replication fork-like structures and D-loops. This may be mediated in part by the OB-fold of MCM8IP, which can exhibit ssDNA-binding activity (Figure 4A). Alternatively, the ability to bind RPA may allow MCM8IP to direct MCM8-9 to RPA-coated ssDNA within functionally relevant substrates. For example, RPA-bound ssDNA on the template strand may facilitate the positioning of MCM8-9 to promote extension of a D-loop^6^.

Another potential function of MCM8IP could be in regulating the enzymatic activities of MCM8-9. MCM8 exhibits DNA-dependent ATPase activity, while both MCM8 and MCM9 display helicase activities *in vitro*^9, 43^. It would therefore be interesting to determine whether physical interaction with MCM8IP directly stimulates these activities. Relatedly, MCM8IP interaction may enhance the processivity of MCM8-9 helicase activity. Previous studies and ours demonstrate that both MCM8-9 and MCM8IP exhibit ssDNA-binding activity (Figure 4A-C)^42^. As a complex, MCM8IP-MCM8-9 may therefore have higher affinity for ssDNA, which potentially allows for increased transactions on DNA due to reduced dissociation. Alternatively, MCM8IP binding may facilitate a structural transition from a MCM8-9 heterodimer to a heterohexamer, thereby enhancing processivity due to encirclement of ssDNA^15, 19^. Lastly, in lieu of enhancing processivity, the RPA molecules that stabilize D-loops may increase the local availability of MCM8-9 through MCM8IP interaction in order to effectively promote continuous D-loop extension and migration.

### MCM8IP and the regulation of replication fork progression and restart after DNA damage

Our findings indicate that MCM8IP and its interactions with RPA and MCM8-9 are required to maintain replication fork progression in the presence of cisplatin (Figure 7A-D). In addition, MCM8IP promotes the restart of stalled forks upon cisplatin treatment (Supplementary Figure 7A). The reasons for the impairment in fork progression and restart in the absence of MCM8IP remain to be clarified but possibilities include persistent interstrand crosslink (ICL)-induced fork reversal and defective replication-coupled ICL traverse or repair.

Upon DNA damage, replication forks undergo reversal, resulting in the annealing of the newly synthesized DNA strands into a fourth fork arm^44^. Fork reversal helps maintain genome stability by restraining fork progression in the presence of DNA damage and allowing sufficient time for the repair of DNA lesions^44^. In response to ICL-inducing agents, fork reversal can occur globally at sites distal to ICL lesions or locally at sites of ICL encounter^45–47^. Fork reversal is mediated by RAD51^46^ or the SNF2 family DNA translocases SMARCAL1, ZRANB3 and HLTF^47–54^ and is controlled globally by ATR signaling in response to ICL-inducing agents^45^. Global fork reversal has been suggested to be a protective mechanism to limit ICL encounters^45^, while local fork reversal facilitates ICL repair or traverse^38, 55–58^. The defective fork progression and restart observed in MCM8IP-deficient cells upon cisplatin treatment could result from persistent global fork reversal due to either a direct or indirect mechanism. Directly, MCM8-9 may be required to physically reset reversed forks in a manner dependent on MCM8IP and its interaction with RPA. Indirectly, elevated ATR signaling in MCM8IP-deficient cells, perhaps due to unrepaired ICLs, may continuously promote fork reversal. In either case, persistent reversed forks may be subjected to cleavage by structure-specific endonucleases, thus requiring MCM8IP-dependent HR to restart the resulting collapsed forks. Further studies are needed to explore these possibilities.

The impairment in fork progression and restart observed in MCM8IP-deficient cells upon cisplatin treatment may also be indicative of defective ICL traverse. ICLs are considered significant barriers to active replication forks that can impede both leading strand synthesis and replicative helicase progression^37, 38^. However, a mechanism involving traverse of an ICL by the replisome without significantly compromising fork progression has recently been described^56^. While the exact details of this process need to be elucidated, ICL traverse involves remodeling of the CDC45-MCM2-7-GINS (CMG) helicase^59^. It has been suggested that FANCM, BLM^60^, or an as yet unidentified helicase(s) facilitates replisome traverse in part by generating ssDNA beyond the ICL, which allows an open replicative helicase complex to re-encircle DNA after translocation^37^. It is possible that the MCM8IP-MCM8-9 complex generates the ssDNA patch required for the loading of the MCM2-7 complex. Alternatively, RPA-mediated stabilization of ssDNA regions generated by another helicase could in turn recruit MCM8-9 through MCM8IP to directly promote repriming of DNA synthesis downstream of the ICL as an alternative to MCM2-7.

Besides ICL traverse, ICL repair can also promote the restart of DNA synthesis when fork progression is arrested by an ICL^37, 38^. ICL repair is mediated by the FA pathway, which coordinates the formation of dual incisions to unhook the crosslink^61–63^. Crosslink unhooking generates DSBs that are repaired by HR to restart collapsed forks^57, 58, 64, 65^. MCM8IP and its interaction with MCM8-9 and RPA may therefore be also required for HR-dependent DNA synthesis during ICL repair.

### MCM8IP as a therapeutic target in cancer

Our work identifies the importance of the MCM8IP-MCM8-9 complex in mediating cellular resistance to cisplatin and olaparib. Interestingly, MCM8IP and MCM8-9 have recently been identified in genetic screens for mediators of ATR inhibitor and temozolomide resistance^66, 67^. Collectively, these studies implicate a broad role for the MCM8IP-MCM8-9 complex in promoting chemoresistance to clinically relevant chemotherapeutic agents.

This raises the possibility of targeting MCM8IP or its interaction with MCM8-9 in combination with DNA damaging agents in the treatment of cancer. One possibility is the use of small molecules that inhibit the ATPase activity of MCM8-9 or disrupt the MCM8IP-MCM8-9 interaction. Small peptides that encompass the MCM8-9 binding motif of MCM8IP (Figure 3E) could also be employed to specifically target the MCM8IP-MCM8-9 interaction.

MCM8-9 have been proposed to mediate RAD51-dependent BIR^19^. This pathway does not promote mitotic DNA synthesis (MiDAS), which occurs through a RAD52-dependent microhomology-mediated BIR (MMBIR) mechanism^68^. Interestingly, while the above BIR pathways share the need for POLD3 for DNA synthesis^69, 70^, they appear mechanistically different. Given that targeting MCM8-9 increases cellular dependency on MiDAS^19^, it might therefore be advantageous to target both BIR pathways in cancer cells in the context of replication stress. As ATR and RAD52 inhibition synergistically cause cell death in the context of replication stress^68^, we propose a strategy of further inhibiting the MCM8IP-MCM8-9 complex in combination with ATR and RAD52 inhibition to exacerbate replication stress in cancer cells.

### MCM8IP and Primary Ovarian Insufficiency

POI is an infertility syndrome caused by mutations in at least 60 genes^71^. Among these genes are *MCM8* and *MCM9*, whose mutations in patient-derived cells cause elevated ICL-induced chromosomal instability^41, 42, 72^. MCM8-9-null mice exhibit infertility due to gametogenesis defects^13, 14^. Collectively, POI may be a consequence of defective MCM8-9-dependent HR repair. Our work implicates the MCM8IP-MCM8-9 complex in HR repair and resistance to ICL-inducing agents, raising the prospect that mutations in MCM8IP, particularly those that disrupt its interaction with MCM8-9, may also cause POI. Reciprocally, mutations in MCM8 or MCM9 that disrupt interactions with MCM8IP may also predispose to POI (Figure 7E). These possibilities provide the rationale for the generation of a mouse model for MCM8IP deficiency to study POI etiology. By providing novel insights into the processes that cause POI, these studies should reveal novel therapeutic opportunities for POI patients.

## METHODS

### Cell culture

HEK293T, HEK293T T-REx, U2OS, U2OS DR-GFP and U2OS 35S (SCR/RFP) cells were cultured in DMEM supplemented with 10% Fetalgro bovine growth serum (RMBIO). HCT116 cells were cultured in RPMI 1640 supplemented with 10% Fetalgro bovine growth serum. Cells were grown in humidified incubators at 37°C and 5% CO_2_.

### Plasmids

pDONR223-RPA1 is previously described^25^. MCM8IP cDNA (C17orf53; NM_001171251) amplified from MDA-MB-436 and MCM8 (NM_032485) and MCM9 (NM_017696) cDNAs amplified from HEK293T were recombined into pDONR223 with BP clonase II (Thermo Fisher). Site-directed mutagenesis by inverse PCR was used to generate mutations in MCM8IP and MCM8. All constructs generated for this study were verified by Sanger sequencing. Gateway destination vectors used in this study include pET59-DEST, pET60-DEST, pMSCV-FLAG-HA-DEST, pHAGE-Ct-FLAG-HA-DEST and the constructs described below. Gateway recombination was performed with LR Clonase II (Thermo Fisher). Gateway destination lentiviral vectors for 5’- and 3’-end tagging with BioID were constructed for this study (pHAGE-TREX-NtBioID-DEST and pHAGE-TREX-CtBioID-DEST). Briefly, BioID was amplified from pcDNA3.1 MCS-BirA(R118G)-HA (Addgene #36047) with a 5’- or 3’-end HA-tag and cloned with a linker (13 x GGGGS) either upstream or downstream of the attP1/2 recombination cassette of pHAGE-TREX-DEST-puro. For complementation studies in MCM8IP knockout cells, a Gateway destination lentiviral vector for UbC promoter-driven transgene expression was constructed (pHAGE-UbC-Hygro-DEST). Briefly, the UbC promoter from pPB-UbC^73^ was cloned into the NheI/BamHI sites of pHAGE-TREX-DEST, replacing the CMV promoter and Tet-responsive elements. The puromycin resistance gene was replaced with hygromycin resistance gene by standard cloning techniques.

### Antibodies

The antibodies used in this study are as follows: rabbit anti-C17orf53/MCM8IP (Sigma HPA023393), mouse anti-C17orf53/MCM8IP (Novus NBP2-37407), rabbit anti-MCM8 (Proteintech 16451-1-AP), rabbit anti-MCM9 (Millipore ABE2603), rabbit anti-RPA1 (Bethyl A300-241A), mouse anti-GST (Santa Cruz sc-138), rabbit anti-GST (Abcam ab21070), mouse anti-FLAG (Sigma F1804), mouse anti-HA (Sigma H3663), rabbit anti-RPA32 (Bethyl A300-244A), rat anti-tubulin (Novus NB600-506), mouse anti-vinculin (Sigma V9131), rabbit anti-SMARCAL1 (Bethyl A301-616A), rabbit anti-RAD51 (BioAcademia BAM-70-002), mouse anti-cyclin A (Santa Cruz sc-271682), rabbit anti-histone H3 (Bethyl A300-823A), rabbit anti-γH2AX (Bethyl A300-081A), rat anti-BrdU (Novus NB500-169), and mouse anti-BrdU (BD Biosciences 347580).

### Protein purification

Human MCM8-9, MCM8IP WT and MBM #1 mutant were expressed in *Spodoptera frugiperda* 9 (*Sf*9) insect cells. MCM9 was cloned into the NotI and SalI sites of pFastBac1-MBP-CtIP-his^74^ to obtain pFastBac1-MBP-MCM9 and MCM8 was cloned into pFastBac1 (Thermo Scientific) using the BamHI and XbaI sites to obtain pFastBac1-FLAG-MCM8. The sequence coding for MCM8 and MCM9 was codon-optimized for expression in *Sf*9 cells (Gen9). Recombinant MCM8-9 was expressed and purified as a complex in *Sf*9 cells by coinfection with baculoviruses prepared from individual pFastBac1 plasmids. MCM8IP WT and MBM #1 were prepared using the same procedure from pFasBac1 plasmids coding for C-terminal FLAG-tagged proteins. Bacmids, primary and secondary baculoviruses for all constructs were prepared using standard procedures according to manufacturer’s instructions (Bac-to-Bac, Life Technologies). The transfection of *Sf*9 cells was carried out using a Trans-IT insect reagent (Mirus Bio).

For large scale expression and purification of MCM8-9, 800 ml of *Sf*9 cells were seeded at 0.5 x 10^6^ per ml and co-infected with 1:1 ratio of both recombinant baculoviruses. The infected cells were incubated in suspension at 27°C for 52 hours with constant agitation. All purification steps were carried out at 4°C or on ice. The *Sf*9 cell pellets were resuspended in 3 volumes of lysis buffer containing 50 mM Tris-HCl (pH 7.5), 1 mM DTT, 1 mM ethylenediaminetetraacetic acid (EDTA), 1:400 protease inhibitory cocktail (Sigma P8340), phenylmethylsulfonyl fluoride (PMSF) and 30 µg/ml leupeptin for 20 min with continuous stirring. Glycerol was added to 16% (v/v) concentration. Next, 5 M NaCl was added slowly to reach a final concentration of 305 mM. The cell suspension was further incubated for 30 min with continuous stirring, centrifuged at 50,000 g for 30 min to obtain soluble extract. Pre-equilibrated amylose resin (New England Biolabs) was added to the cleared extract and incubated for 1 hour with continuous mixing. The amylose resin was separated from the soluble extract by centrifugation at 2000 g for 2 min and the supernatant was discarded. The amylose resin was washed extensively batch-wise as well as on disposable columns (Thermo Scientific) with wash buffer (50 mM Tris-HCl pH 7.5, 2 mM β-mercaptoethanol, 300 mM NaCl, 10 % glycerol, 1 mM PMSF). The bound proteins were eluted with elution buffer (50 mM Tris-HCl pH 7.5, 0.5 mM β-mercaptoethanol, 300 mM NaCl, 10 % glycerol, 1 mM PMSF, 10 mM maltose). The eluate was then treated with 1/10 (w/w) PreScission protease for 60 min to cleave off the maltose binding protein (MBP) affinity tag. The cleaved sample was added to pre-equilibrated anti-FLAG M2 Affinity Gel (A2220, Sigma) for 1 hour with continuous mixing. The FLAG resin was washed extensively on a disposable column (Thermo Scientific) with FLAG wash buffer (50 mM Tris-HCl pH 7.5, 150 mM NaCl, 10 % glycerol, 1 mM PMSF, 0.5 mM β-mercaptoethanol). Finally, recombinant MCM8-9 was eluted from the FLAG resin by FLAG wash buffer supplemented with FLAG peptide (200 ng/µl, Sigma, F4799) and stored at – 80°C.

For large-scale expression and purification of MCM8IP WT and mutant, 1000 ml of *Sf*9 cells were used and the soluble extracts were prepared as described above. Pre-equilibrated anti-FLAG M2 Affinity Gel (A2220, Sigma) was added to each of the soluble extract for 1 hour with continuous mixing. The FLAG resin was separated and washed batch wise as described above. The FLAG resin was then washed extensively on a disposable column (Thermo Scientific) with FLAG wash buffer (50 mM Tris-HCl pH 7.5, 300 mM NaCl, 10 % glycerol, 1 mM PMSF, 0.5 mM β-mercaptoethanol, 0.1 % NP 40) and then with FLAG wash buffer containing 100 mM NaCl. Finally, the recombinant protein was eluted from the FLAG resin by FLAG wash buffer (with 100 mM NaCl) supplemented with FLAG peptide (200 ng/µl, Sigma, F4799) and stored at –80°C.

### *In vitro* interaction studies

Overnight cultures of *E. coli* BL21 (DE3) cells carrying pET60-MCM8IP and its mutants or pET59-MCM8 were diluted into 50 ml of LB to OD_600_=0.075 and grown at 30°C to OD_600_=0.5-0.6 (approximately 3 hours). Cultures were then induced with IPTG (1 mM) and grown at 30°C for an additional 6 hours. E. coli BL21 (DE3) carrying pCDFDuet-RPA1 ^25^ were grown overnight at room temperature following induction with IPTG. Cell pellets were lysed in 1 ml wash buffer (20 mM Tris-HCl pH 7.5, 200 mM NaCl, 0.5% NP40, 1 mM DTT, 10% glycerol) supplemented with a protease inhibitor cocktail (Goldbio GB-330), PMSF (1 mM), lysozyme (0.4 mg/ml), DNase I (20 units, New England Biolabs M0303) and MgCl_2_ (10 mM). Lysates were sonicated (3 x 10 sec pulses at 30% output) and then cleared by centrifugation. Cleared lysates from bacteria expressing pET60-MCM8IP and its mutants were incubated with glutathione-agarose beads (Goldbio, G-250) for 2 hours at 4°C with gentle agitation. Immobilized proteins were then washed 4 times and incubated with cleared lysates from bacteria expressing pCDFDuet-RPA1 or pET59-MCM8 for 2 hours at 4°C with gentle agitation. Immobilized protein-complexes were then washed five times, eluted with LDS sample buffer and resolved by SDS-PAGE.

To study the interaction between recombinant MCM8-9 and MCM8IP, *Sf*9 cells were infected with MBP-MCM9 and FLAG-MCM8 baculoviruses (see Protein Purification). Cells were lysed and MCM8-9 was immobilized on amylose resin (New England Biolabs) and washed with wash buffer (50 mM Tris-HCl pH 7.5, 2 mM β-mercaptoethanol, 300 mM NaCl, 0.1% (v/v) NP40, 1 mM PMSF). Resin-bound MCM8-9 was then incubated with 1 µg of MCM8IP, either wild-type or MBM #1 mutant, diluted in binding buffer (50 mM Tris-HCl pH 7.5, 2 mM β-mercaptoethanol, 3 mM EDTA, 100 mM NaCl, 0.2 µg/µl BSA, 1 mM PMSF) for 1 hour at 4°C with continuous rotation. The resin was washed four times with wash buffer containing 100 mM NaCl, proteins were eluted in wash buffer supplemented with 10 mM maltose and detected by western blotting. As a negative control, MCM8IP was incubated with the resin without the bait protein.

### Immunoprecipitation

Immunoprecipitation was performed as previously reported^25^. Briefly, HEK293T cells transduced with retroviruses carrying pMSCV-FLAG-HA-RPA1, -MCM8IP, -MCM8, - MCM9 or a -GFP control were grown to near confluency in a 10 cm dish and harvested in PBS. Cell pellets were resuspended in 750 µl of MCLB buffer (50 mM Tris-HCl pH 7.5, 1% NP40) supplemented with 150 mM NaCl, and protease and phosphatase inhibitor cocktails (Goldbio, GB-331 and GB-450). Following incubation for 30 min at 4°C with gentle agitation, cell lysates were cleared by centrifugation and the low-salt supernatant collected. Cell pellets were then resuspended in 250 µl of MCLB supplemented with 500 mM NaCl and protease and phosphatase inhibitors and gently agitated for 1 hour at 4°C. After centrifugation, the salt concentration of the supernatant was adjusted to 150 mM NaCl and combined with the low-salt supernatant. The combined lysates were then incubated with 20 µl of anti-HA agarose beads (Sigma-Aldrich A2095) for 4 hours at 4°C with gentle agitation. Protein-bound beads were subsequently washed four times in buffer (50 mM Tris-HCl pH 7.5, 1% NP-40 and 150 mM NaCl) and bound proteins eluted in LDS sample buffer. For mass spectrometry, HEK293T cells stably transduced with pHAGE-MCM8IP-FLAG-HA or pHAGE-GFP-FLAG-HA were grown to near confluency in two 15 cm dishes and the method described above was scaled up accordingly.

### Proximity-dependent labeling with BioID

Small-scale BioID experiments were performed with cells grown to near confluency in 10 cm dishes. Briefly, HEK293T T-REx cells transduced with lentivirus carrying pHAGE-TREX-BioID control or pHAGE-TREX-BioID-RPA1 were treated with 1 µg/ml doxycycline for 24 hours. Cells were then treated with DNA damaging agents or left untreated in media supplemented with 50 µM biotin and 1 µg/ml doxycycline for an additional 18-24 hours. Cells were washed 3 times and harvested in PBS. Cell pellets were resuspended in 1 ml of RIPA buffer (50 mM Tris-HCl pH 7.5, 150 mM NaCl, 1% NP40, 0.5% deoxycholic acid, 0.1% SDS, 1 mM EDTA, 1 mM DTT) supplemented with protease and phosphatase inhibitors. Lysates were then sonicated (3 x 10 sec pulses at 30% output), and treated with benzonase for 30 min at 4°C with gentle agitation. Following centrifugation, cleared lysates were incubated with 20 µl streptavidin-coated magnetic beads (Thermo Fisher 65001) for 4 hours at 4°C with gentle agitation. Protein-bound beads were separated using a magnetic rack, washed 4 times with RIPA buffer, and eluted with LDS sample buffer supplemented with biotin.

To identify RPA1 interactors by mass spectrometry, HEK293T T-REx cells expressing pHAGE-TREX-BioID-RPA1 or the BioID-alone control were grown to near confluency in three 15 cm dishes and labeled and harvested as described above. Cells from each sample were then evenly distributed among seven microfuge tubes. Each cell pellet was resuspended in 1 ml of lysis buffer (6 M Urea, 50 mM Tris-HCl pH 7.5, 0.5% Triton X-100) supplemented with 1 mM DTT and protease inhibitors, sonicated (3 x 10 sec pulses at 30% output) and cleared by centrifugation. Lysates were then incubated with 40 µl of streptavidin-coated magnetic beads per tube (280 µl total beads per sample) overnight at 4°C with gentle agitation. The next day, protein-bound beads for each sample were pooled and washed 4 times with lysis buffer and another 4 times with lysis buffer lacking Triton X-100 while exchanging fresh microfuge tubes with each wash. All remaining buffer from the final wash was removed by aspiration.

To identify MCM8IP interactors by mass spectrometry, HEK293T T-REx cells expressing pHAGE-TREX-BioID-MCM8IP, pHAGE-TREX-MCM8IP-BioID or the BioID-alone control were grown to near confluency in three 15 cm dishes and labeled and harvested as above. Following the even distribution of cells into seven microfuge tubes, each cell pellet was resuspended in 1 ml RIPA Buffer and lysates were prepared as described above for small-scale purification. Prepared lysates were incubated with 20 µl streptavidin beads per tube (140 µl total beads per sample) for 4 hours at 4°C with gentle agitation. Protein-bound beads were pooled and subsequently washed 4 times with RIPA buffer and another 4 times with detergent-free buffer (50 mM Tris-HCl pH 7.5, 150 mM NaCl) while exchanging fresh microfuge tubes with each wash. All remaining buffer from the final wash was removed by aspiration.

### In-gel digestion of immunoprecipitated samples for mass spectrometry

Immunoprecipitated samples were separated on 4-12% gradient SDS-PAGE, and stained with SimplyBlue (Thermo Fisher). Protein gel slices were excised and *in-gel* digestion performed. Gel slices were washed with 1:1 (Acetonitrile: 100 mM ammonium bicarbonate) for 30 min and then dehydrated with 100% acetonitrile for 10 min until shrinkage. Excess acetonitrile was removed and slices were dried in a speed-vac for 10 min without heat. Gel slices were then reduced with 5 mM DTT for 30 min at 56°C in an air thermostat and chilled down to room temperature before alkylation with 11 mM IAA for 30 min in the dark. Gel slices were washed with 100 mM ammonium bicarbonate and 100 % acetonitrile for 10 min each. Excess acetonitrile was removed and dried in a speed-vac for 10 min at no heat. Gel slices were then rehydrated in a solution of 25 ng/µl trypsin in 50 mM ammonium bicarbonate on ice for 30 min. Digestions were performed overnight at 37°C in an air thermostat. Digested peptides were collected and further extracted from gel slices in extraction buffer (1:2 v/v, 5% formic acid/acetonitrile) with high-speed shaking in an air thermostat. Supernatant from both extractions were combined and dried down in a speed-vac. Peptides were dissolved in 3% acetonitrile/0.1% formic acid.

### On-bead digestion of BioID samples for mass spectrometry

Proteins bound to streptavidin beads were washed five times with 200 µl of 50 mM ammonium bicarbonate and subjected to disulfide bond reduction with 5 mM DTT (56°C, 30 min) and alkylation with 10 mM iodoacetamide (room temperature, 30 min in the dark). Excess iodoacetamide was quenched with 5 mM DTT (room temperature, 15 min in the dark). Proteins bound on beads were digested overnight at 37°C with 1 µg of trypsin/LysC mix. The next day, digested peptides were collected in a new microfuge tube and digestion was stopped by the addition of 1% TFA (final v/v), and centrifuged at 14,000 g for 10 min at room temperature. Cleared digested peptides were desalted on a SDB-RP StageTip and dried in a speed-vac. Dried peptides were dissolved in 3% acetonitrile/0.1% formic acid.

### LC-MS/MS analysis

Thermo Scientific UltiMate 3000 RSLCnano system, Thermo Scientific EASY Spray source with Thermo Scientific Acclaim PepMap100 2 cm x 75 µm trap column, and Thermo Scientific EASY-Spray PepMap RSLC C18 were used for peptide preparation. 50 cm x 75 µm ID column were used to separate desalted peptides with a 5-30% acetonitrile gradient in 0.1% formic acid over 127 min at a flow rate of 250 nl/min. After each gradient, the column was washed with 90% buffer B (0.1% formic acid, 100% HPLC-grade acetonitrile) for 5 min and re-equilibrated with 98% buffer A (0.1% formic acid, 100% HPLC-grade water) for 40 min.

Thermo Scientific Q Exactive HF mass spectrometer was used for peptide MS/MS analysis of BirA*-RPA1 and BirA* control. MS data were acquired with an automatic switch between a full scan and 15 data-dependent MS/MS scans (TopN method). Target value for the full scan MS spectra was 3 x 10^6^ ions in the 375-2000 m/z range with a maximum injection time of 100 ms and resolution of 60,000 at 200 *m/z* with data collected in profile mode. Precursors were selected using a 1.6 *m/z* isolation width. Precursors were fragmented by higher-energy C-trap dissociation (HCD) with normalized collision energy of 27 eV. MS/MS scans were acquired at a resolution of 15,000 at 200 m/z with an ion target value of 2 x 10^5^, maximum injection time of 50 ms, dynamic exclusion for 15 s and data collected in centroid mode.

Thermo Scientific Orbitrap Fusion Tribrid mass spectrometer was used for peptide MS/MS analysis of BirA*-MCM8IP, MCM8IP-BirA* and MCM8IP-HA and respective controls. Survey scans of peptide precursors were performed from 400 to 1575 *m/z* at 120K FWHM resolution (at 200 *m/z*) with a 2 x 10^5^ ion count target and a maximum injection time of 50 msec. The instrument was set to run in top speed mode with 3 sec cycles for the survey and the MS/MS scans. After a survey scan, tandem MS was performed on the most abundant precursors exhibiting a charge state from 2 to 6 of greater than 5 x 10^3^ intensity by isolating them in the quadrupole at 1.6 Th. CID fragmentation was applied with 35% collision energy and resulting fragments were detected using the rapid scan rate in the ion trap. The AGC target for MS/MS was set to 1 x 10^4^ and the maximum injection time limited to 35 msec. The dynamic exclusion was set to 45 sec with a 10 ppm mass tolerance around the precursor and its isotopes. Monoisotopic precursor selection was enabled.

### Mass spectrometry data analysis

Raw mass spectrometric data were processed and searched using the Sequest HT search engine within the Proteome Discoverer 2.2 (PD2.2, Thermo Fisher) with custom sequences and the reference *Saccharomyces cerevisiae* database downloaded from Uniprot. The default search settings used for protein identification in PD2.2 searching software were as follows: two mis-cleavages for full trypsin with fixed carbamidomethyl modification of cysteine and oxidation of methionine and deamidation of asparagine and glutamine and acetylation on N-terminal of protein were used as variable modifications. Identified peptides were filtered for maximum 1% (BirA*-RPA1) or 5% (BirA*-MCM8IP, MCM8IP-BirA*, MCM8IP-HA) false discovery rate using the Percolator algorithm in PD2.2. PD2.2 output combined folder was uploaded in Scaffold (Proteome Software) for data visualization. Spectral counting was used for analysis to compare the samples.

### Electrophoretic mobility shift assays

Single stranded DNA oligonucleotide (93 nt long, X12-3HJ3)^75^ was labeled at the 5’ terminus with [γ-^32^P] ATP and T4 polynucleotide kinase (New England Biolabs), according to standard protocols. Unincorporated nucleotides were removed using Micro Bio-Spin P-30 Gel Columns (Bio-Rad). The binding reactions (15 µl volume) were carried out in 25 mM Tris acetate, pH 7.5, 3 mM EDTA, 1 mM DTT, 100 µg/ml BSA (New England Biolabs), single stranded DNA substrate (1 nM, molecules). Proteins were added and incubated for 15 min on ice. Loading dye (5 µl; 50% glycerol, bromophenol blue) was added to reactions and products were separated on polyacrylamide gels (ratio acrylamide:bisacrylamide 19:1, Bio-Rad) in TBE buffer at 4°C. The gels were dried on 17 CHR (Whatman), exposed to a storage phosphor screen (GE Healthcare) and scanned by a Typhoon Phosphor Imager (FLA9500, GE Healthcare).

### Immunofluorescence

For UV laser microirradiation experiments, U2OS cells expressing pMSCV-FLAG-HA-MCM8IP or pHAGE-Ct-FLAG-HA-MCM8IP and mutants were seeded at a density of 30,000 cells per well in 8-well chamber slides (Nunc). Cells were pre-sensitized with 10 µM BrdU for at least 24 hours. UV laser microirradiation was performed using an Inverted Zeiss AxioObserver.Z1 with PALM MicroBeam IV (cutting parameters: focus 65%, energy 47%, Speed 77%). Following microirradiation, cells were incubated for 2-3 hours at 37°C and then simultaneously fixed and permeabilized (2% formaldehyde, 0.5% Triton X-100 in PBS) for 25 min. Cells were incubated with anti-FLAG (1:1000) and anti-γH2AX (1:1000) overnight in blocking buffer (3% BSA, 0.05% Triton-X100 in PBS). Cells were then stained with appropriate secondary antibodies conjugated with Alexa 488 or Alexa 594 and mounted with Fluoroshield with DAPI. Cells were imaged with a Nikon Eclipse 90i microscope equipped with an industrial camera (The Imaging Source DMK 33UX174) and operated with Nikon NIS Elements software. Images were analyzed in ImageJ.

For RAD51 immunofluorescence studies, HCT116 cells were seeded on black 96-well bottom-glass plates and the day after treated or not with 10 µM cisplatin or subjected to ionizing radiation (8 Gy). Six hours after irradiation or 24 hours after cisplatin treatment, cells were simultaneously fixed and permeabilized (4% paraformaldehyde, 0.5% Triton X-100) for 10 min at room temperature. Cells were incubated in blocking solution (3% bovine serum albumin in TBS-Tween 0.1%) for 1 hour and then in primary antibody diluted in blocking solution for 1 hour at room temperature or overnight at 4°C. Primary anti-RAD51 antibody (Bioacademia 70-002) was used at 1:10,000 dilution and anti-cyclin A antibody (Santa Cruz sc-271682) was used at 1:500 dilution. Cells were washed 3 times with TBS-T and then incubated for 1 hour at room temperature with the appropriate secondary antibody, Alexa Fluor 488-labeled anti-rabbit and Alexa Fluor 594-labeled goat anti-mouse at 1:1000 dilution (Thermo Fisher A-11008 and A-11005). After three washes in TBS-T, cells were incubated with DAPI for 5 min at room temperature to counterstain nuclei. Two-dimensional acquisitions were made using the ImageXpress Nano Automated Imaging System microscope (Molecular Devices) equipped with a 40× Plan Apo objective (0.95 numerical aperture). An integrated imaging software (MetaXpress) was used for image analysis. The total number of RAD51 foci per cell was measured in cyclin A-positive and -negative cells. At least 1000 cells per experimental point were counted, and each experiment was repeated at least 2 times independently.

### Subcellular fractionation

Subcellular fractionation was performed as previously described with some modifications^76^. Briefly, cells were harvested by trypsinization, washed with PBS, and resuspended in CSK buffer (10 mM PIPES pH 6.8, 100 mM NaCl, 1 mM EGTA, 1 mM EDTA, 300 mM sucrose, 1.5 mM MgCl_2_, 0.1% Triton X-100, 1 mM DTT) supplemented with protease and phosphatase inhibitors. After 5 min of incubation on ice, soluble and insoluble fractions were separated by centrifugation (1500 g, 5 min, 4°C). The supernatant (soluble fraction) was collected and the pellet was washed once with CSK buffer. After centrifugation, the supernatant was removed and the pellet (chromatin fraction) was resuspended in LDS sample buffer.

### CRISPR-Cas9 gene targeting

Guide RNAs targeting MCM8IP, MCM8 and MCM9 were designed using GPP sgRNA Designer (Broad Institute). The targeted sequences are as follows: MCM8IP #1 (5’-CGACCCCCCTTGAGACCTGGT-3’), MCM8IP #2 (5’-TTCAGTATTGGCTAAAAAAGC-3’), MCM8IP #3 (5’-CAGCTGGATTGGCAATCAGAG-3’), MCM8 #1 (5’-AATTCTATTAGTAATAGCAA-3’), MCM8 #2 (5’-GTGTGTCGAGGCAGGTCATT-3’), MCM9 #1 (5’-ACGGGATTGTAATGCAACGG-3’), and MCM9 #2 (5’-ACACTGTCTGATGTGGGCAA-3’). MCM8IP sgRNAs were cloned into the BsmBI/Esp3I sites of pXPR206 (Addgene #96920). MCM8 and MCM9 sgRNAs were cloned into the BsmBI/Esp3I sites of pLentiCRISPR v2 Blast (Addgene #98293). Following stable lentiviral transduction of cells, targeting efficiencies were evaluated by western blotting.

### Homologous recombination assays

U2OS DR-GFP^31^ or U2OS 35S (SCR/RFP)^35^ cells (gift from Ralph Scully) were seeded at a density of 250,000 cells per well in 6-well plates. The following day, cells were transfected (Mirus LT-1) with 2.5 µg of an I-SceI-expression vector or an empty vector control (gifts from Shan Zha). Parallel transfections with pEGFP-N3 (gift from Jean Gautier) were performed to assess transfection efficiency. Two days after transfection, cells were harvested by trypsinization and resuspended in PBS in preparation for flow cytometry. Approximately 20,000 DR-GFP cells and 200,000 SCR/RFP cells were analyzed per sample, respectively. GFP and/or RFP-positive populations determined by flow cytometry were normalized for transfection efficiency. Gene conversion events are presented as repair efficiencies relative to the non-targeting control.

For experiments using the BFP reporter, BFP-positive HEK293T cells were seeded at 50–70% confluency into 24-well plates and transfected by mixing TransIT-293T (3 µl; Mirus) and 250 ng plasmid pX330 containing a BFP targeting sgRNA along with a plasmid HDR donor (500 ng). The cells were collected 3 days after transfection and analyzed by flow cytometry for GFP-positive cells.

### Survival assays

Survival assays were performed as previously reported^77^. Briefly, HCT116 cells were seeded at a density of 5,000 cells per well in 12-well plates. The next day, cells were treated with DNA damaging agents and allowed to grow for an additional 5 days. Following fixation (10% methanol and 10% acetic acid in water) and staining with crystal violet (1% w/v in methanol), plates were washed thoroughly and allowed to fully dry. Cells were subsequently destained (0.1% w/v SDS in methanol) and the resuspended solution transferred to a 96-well plate for quantification with a spectrophotometer (λ=595 nm). Following background subtraction, cell survival is presented as a percentage of the untreated control.

### DNA fiber analysis

To measure fork elongation rate, exponentially growing HCT116 cells were pulse-labeled with 25 µM CldU (5 min), washed in warm 1X PBS and exposed to 125 µM IdU with or without 30 µM cisplatin (40 min). Alternatively, to measure fork stalling, cells were pulse-labeled with 25 µM CldU (5 min), exposed to 30 µM cisplatin for 1 hour and 35 min in presence of CldU, washed in warm 1X PBS and exposed to 125 µM IdU (40 min). Labeled cells were trypsinized and resuspended in ice-cold PBS at 2 × 10^5^ cells/ml. Two microliters of this suspension were spotted onto a pre-cleaned glass slide and lysed with 10 µl of spreading buffer (0.5% SDS in 200 mM Tris-HCl pH 7.4 and 50 mM EDTA). After 6 min, the slides were tilted at 15° relative to horizontal, allowing the DNA to spread. Slides were air-dried, fixed in methanol and acetic acid (3:1) for 2 min, rehydrated in PBS for 10 min and denatured with 2.5 M HCl for 1 hour at room temperature. Slides were then rinsed in PBS and blocked in PBS + 0.1% Triton X-100 (PBS-T) + 3% BSA for 1 hour at room temperature. Rat anti-BrdU (1:100, Abcam) and mouse anti-BrdU (1:100, Becton Dickinson) were then applied to detect CldU and IdU, respectively. After a 2-hour incubation, slides were washed in PBS and stained with Alexa Fluor 488-labeled goat anti-mouse IgG1 antibody and Alexa Fluor 594-labeled goat anti-rat antibody (1:300 each, Thermo Fisher). Slides were mounted in Prolong Gold Antifade (Thermo Fisher) and stored at −20°C. Replication tracks were imaged on a Nikon Eclipse 50i microscope fitted with a PL Apo 40X/0.95 numerical aperture (NA) objective and measured using ImageJ software. In each experiment, 100 or more individual tracks were measured for fork elongation rate estimation, more than 400 individual tracks were analyzed for fork stalling estimation. Each experiment was repeated at least 2 times independently.

### Phylogenetic analyses

We located *MCM8IP* gene orthologs in distant species using BLASTp, and in difficult cases PSI-BLAST, starting with either the full protein sequence from human, or a truncated human conserved domain spanning amino acid positions 381 – 575. We targeted species’ genomes that were assembled to high quality within the major branches of the eukaryote tree of life. We searched all major opisthokont lineages that had such high-quality nuclear genome sequences and also included the closely related ameobozoa and green plant groups. When no sequence was found in a high-quality genome sequence, the entire taxonomic group was also searched via Taxonomic group filtering available on the BLAST hosted at NCBI. Attempts to find *MCM8IP* in more distant taxonomic groups were not successful.

To test for correlated gene loss, we compared two nested likelihood models of binary trait evolution (independent and dependent) implemented in *BayesTraits*^78, 79^. The “independent” model had 2 free parameters to estimate, the loss rate of gene 1 and the loss rate of gene 2. We set the parameters for gain of traits (alphas) to zero with the “Restrict alpha1 alpha2 0” command, since it is not possible to gain one of these genes given how we define the *MCM8*, *MCM9* and *MCM8IP* genes as being inherited strictly as orthologs. The “dependent” model then divides the 2 loss rate parameters into loss in the presence or absence of the other gene, resulting in 4 free parameters. Rates corresponding to gene gain were set to zero with the “Restrict q12 q13 q24 q34 0” command.

Each model was provided with a phylogenetic tree reflecting the speciation patterns leading to the studied species. The tree was downloaded from TimeTree and matches the major relationships published for these groups^80, 81^. Gene presence/absence data were provided to the models for each pair of genes separately and the models were optimized with 100,000 maximum likelihood tries to ensure model convergence. Maximum likelihood values of the 2 nested models were compared with a likelihood ratio test and the p-value was estimated with the chi-square distribution with 2 degrees of freedom. A low p-value was taken as evidence of rejection of the independent model in favor of the dependent model for that gene pair.

### Coevolutionary analyses

Evolutionary rate covariation (ERC) values were calculated for protein pairs as the correlation coefficient of branch-specific evolutionary rates using protein sequences from 33 mammalian species^82^. ERC between MCM8IP and major mammalian DNA repair pathways was calculated by taking the mean ERC score between MCM8IP and the genes in each pathway. Statistical significance was determined by permutation test, where the p-value is the computed probability of the observed mean ERC or greater when taking the mean between each gene group and 10,000 random genes.

## DATA AVAILABILITY

All unique reagents generated in this study will be made available upon request.

## ACKNOWLEDGMENTS

We thank Richard Baer for critical reading of the manuscript. The U2OS 35S (SCR/RFP) cells were kindly provided by Ralph Scully (Harvard Medical School). This work was supported by the NIH grants R01 CA197774 and R01 GG014860, and the Irma T. Hirschl Award to A.C., NIH T32CA009503 to J.W.H., and the Italian Association for Cancer Research (AIRC) post-doctoral research fellowship to G.L. K.A.B. is supported by NIH grant R01 ES024872, American Cancer Society 129182-RSG-16-043-01-DMC, and Stand Up to Cancer Innovative Research Grant SU2C-AACR-IRG-02-16. N.L.C. is supported by NIH grant R01HG009299. Studies conducted in the Proteomics, Confocal and Specialized Microscopy and Flow Cytometry Shared Resources of the Herbert Irving Comprehensive Cancer Center at Columbia University were supported by the NIH grant P30 CA013696.

## AUTHOR CONTRIBUTIONS

J.W.H. and A.C. conceived the study. J.W.H. identified MCM8IP, conducted all the physical and genetic interaction studies and generated the reagents utilized in this study. A.T. performed DNA replication studies, G.L. conducted genome instability assays, and T.S.N. performed CRISPR-based HR assays. R.C.M. and S.B.H. helped generate KO cells. A.A. and R.A. expressed and purified recombinant MCM8IP and MCM8-9 and performed *in vitro* biochemical assays under the supervision of P.C. G.J.B. performed co-evolutionary analyses under the supervision of K.A.B. N.L.C. performed phylogenetic studies. R.K.S. analyzed the proteomics data. J.W.H. and A.C. wrote the manuscript. All authors read and approved the manuscript.

## SUPPLEMENTARY FIGURE LEGENDS

**Supplementary Figure 1.**
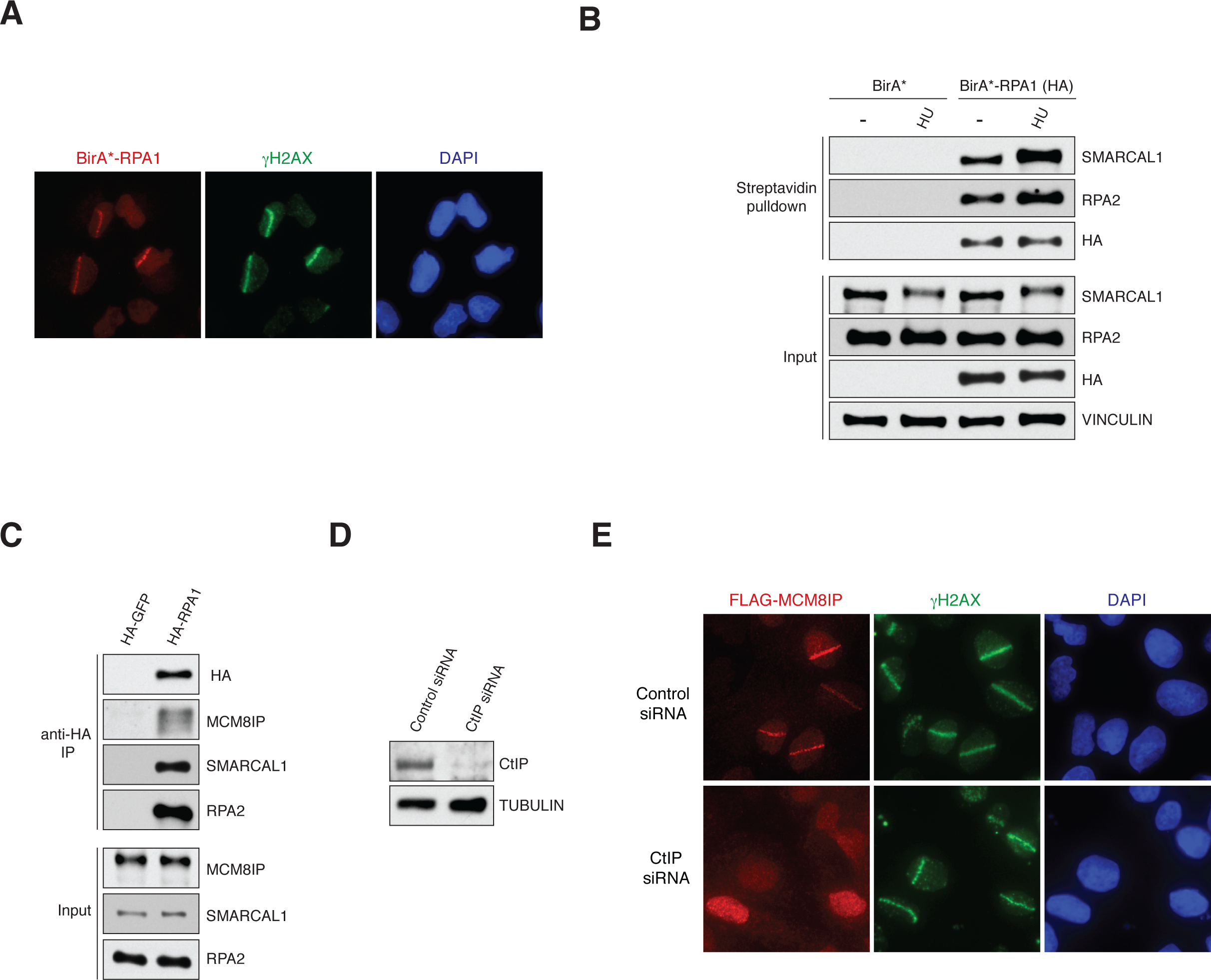
Interaction and localization studies for RPA1 and MCM8IP. (A) Representative images of HA-BirA*-RPA1 recruitment to sites of UV laser microirradiation in U2OS cells. DNA damage tracts are indicated with γH2AX staining. (B) Detection by western blot of RPA2 and SMARCAL1 in streptavidin pulldowns from HEK293T cells expressing BirA* or BirA*-RPA1. Cells were treated with HU (1 mM) in the presence of exogenous biotin for 24 hours prior to lysis. Vinculin is shown as a loading control. (C) Detection by western blot of MCM8IP, SMARCAL1 and RPA2 co-immunoprecipitated by HA-GFP or HA-RPA1 from HEK293T cells. (D) Detection by western blot of CtIP in U2OS cells expressing FLAG-MCM8IP transfected with control or CtIP siRNA. Tubulin is shown as a loading control. (E) Representative images of FLAG-MCM8IP recruitment to sites of UV laser microirradiation in U2OS cells transfected with control or CtIP siRNA. DNA damage tracts are indicated with γH2AX staining.

**Supplementary Figure 2.**
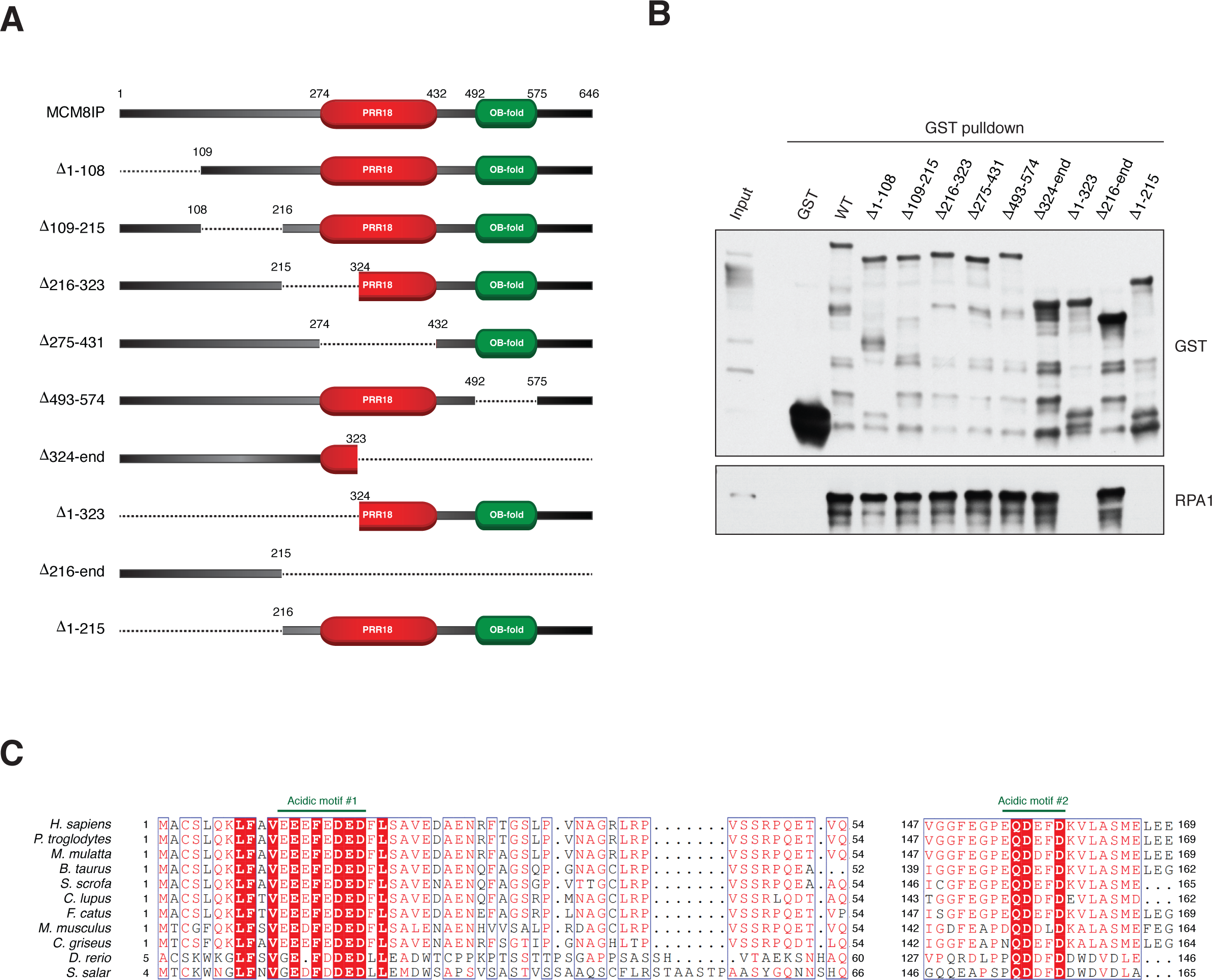
Identification of RPA1-binding motifs in MCM8IP. (A) Schematic representation of full-length MCM8IP and MCM8IP deletion mutants. Shown in red and green are the PRR18 and DUF4539 (a predicted OB-fold) domains, respectively. (B) Detection by western blot of RPA1 co-precipitated by bead-bound recombinant GST, GST-MCM8IP WT or GST fused to MCM8IP mutants presented in (A). (C) Alignment from various species of two conserved acidic motifs within the first 215 amino acids of human MCM8IP predicted to interact with RPA1. Sequence alignments were conducted using Clustal Omega and processed using ESPript.

**Supplementary Figure 3.**
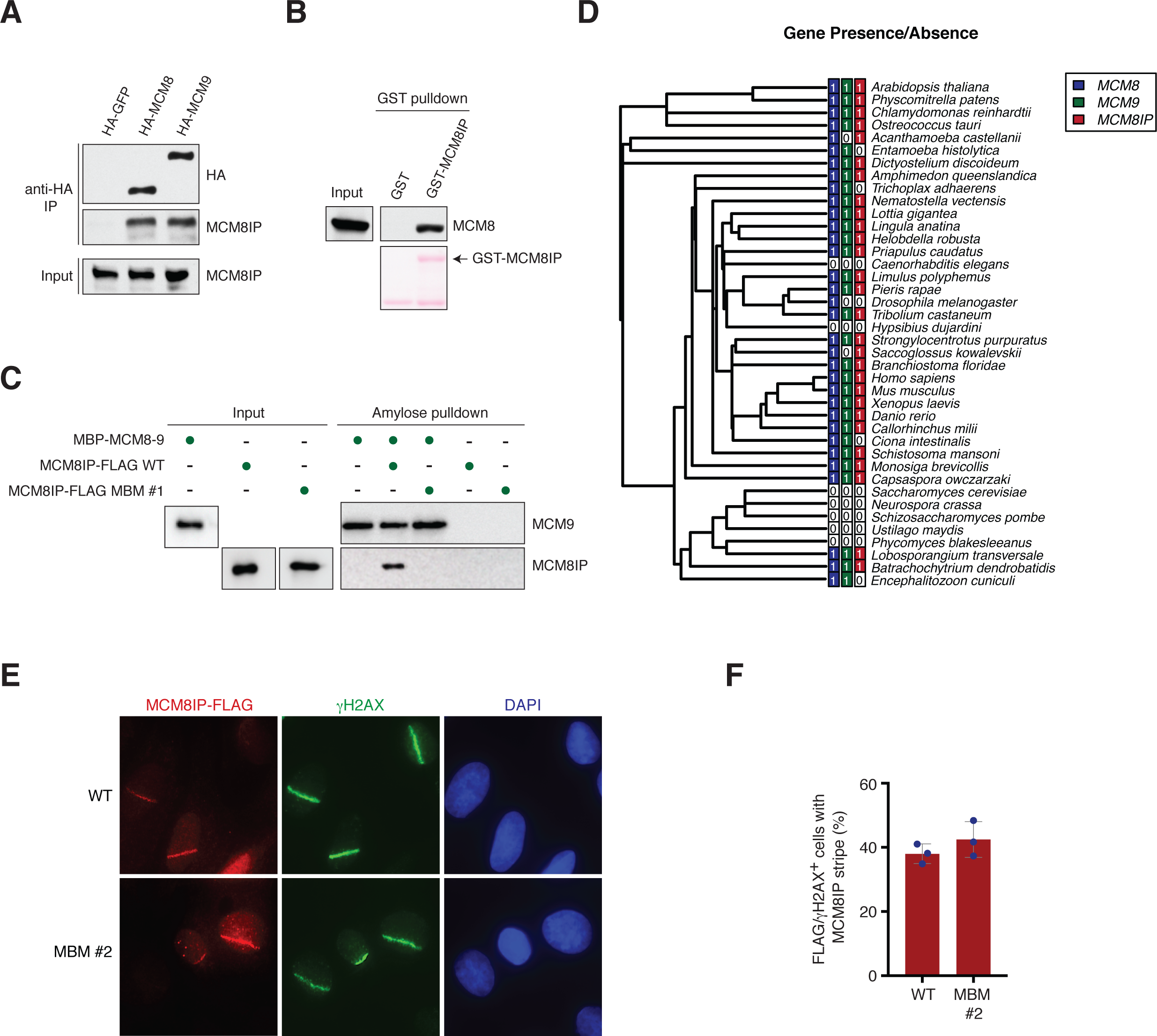
Characterization of the MCM8IP-MCM8-9 interaction. (A) Detection by western blot of MCM8IP co-immunoprecipitated with HA-GFP, HA-MCM8 or HA-MCM9 from HEK293T cells. (B) Detection by western blot of recombinant HIS-MCM8 co-precipitated by bead-bound recombinant GST or GST-MCM8IP. (C) Detection by western blot of recombinant MCM8IP-FLAG, either WT or MBM #1 mutant, co-precipitated by amylose bead-bound recombinant MBP-MCM8-9 complex. (D) Phylogenetic analysis indicating the presence or absence of *MCM8IP*, *MCM8* and *MCM9* in various species. This species phylogeny represents most major taxonomic groups of opisthokonts, amoebas, and green plants. Color-filled squares containing “1” indicate the presence of the gene in the indicated species. (E) Representative images of the recruitment of MCM8IP-FLAG WT or MBM #2 in U2OS cells following UV laser microirradiation. DNA damage tracts are indicated with γH2AX staining. (F) Graphical representation of the percentage of MCM8IP-FLAG WT or MBM #2 co-localizing with γH2AX following UV laser microirradiation in U2OS cells. The mean ± SD of three independent experiments is presented.

**Supplementary Figure 4.**
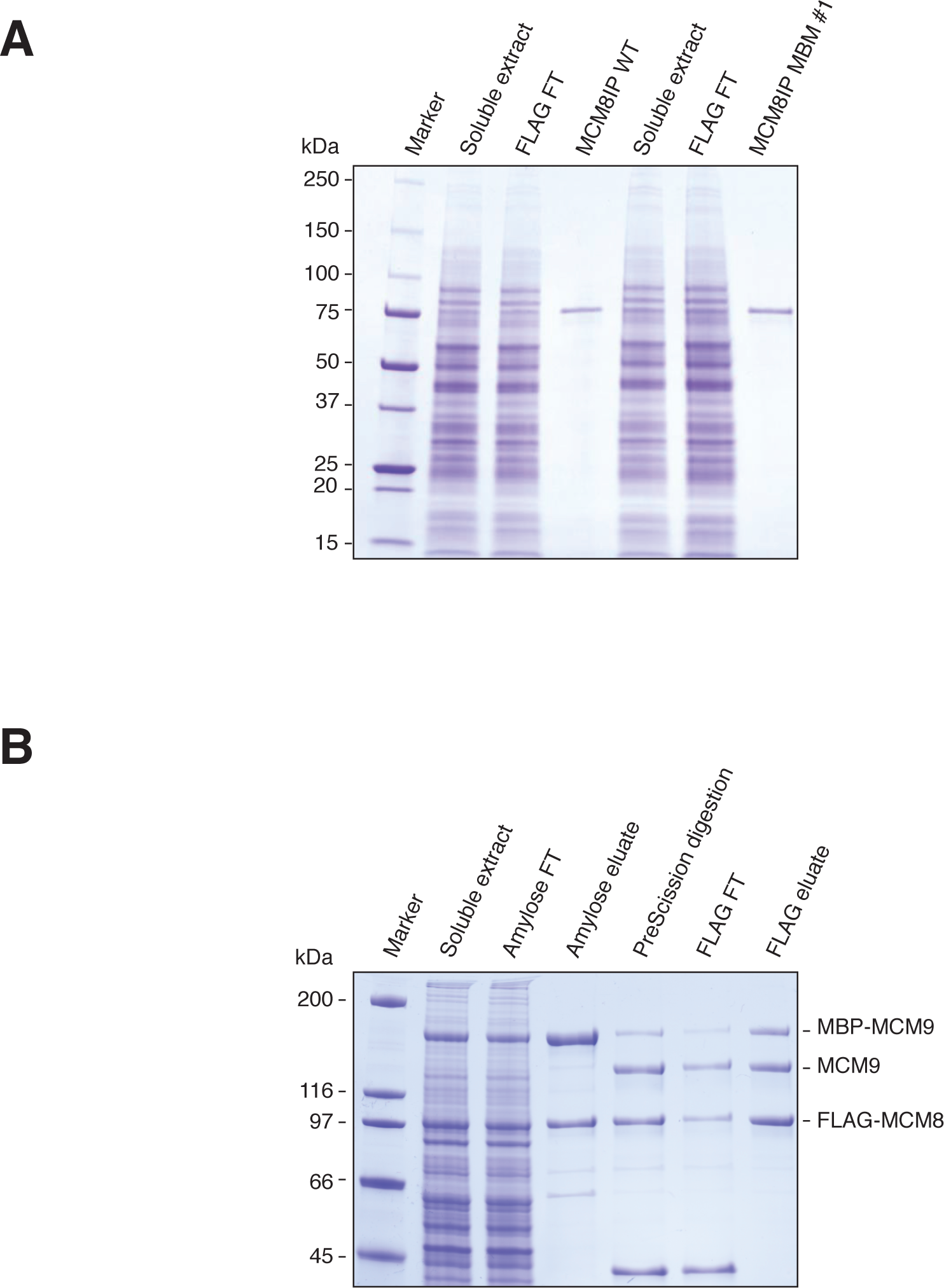
Purification of recombinant MCM8IP and MCM8-9. (A) Coomassie-stained gel showing the purification of recombinant MCM8IP-FLAG WT or MBM #1 from *Sf*9 cells. FLAG FT, flowthrough from the FLAG resin. (B) Coomassie-stained gel showing the steps of the co-purification of recombinant MBP-MCM9 and FLAG-MCM8 from *Sf*9 cells. Amylose FT, flowthrough from the amylose resin; FLAG FT, flowthrough from the FLAG resin.

**Supplementary Figure 5.**
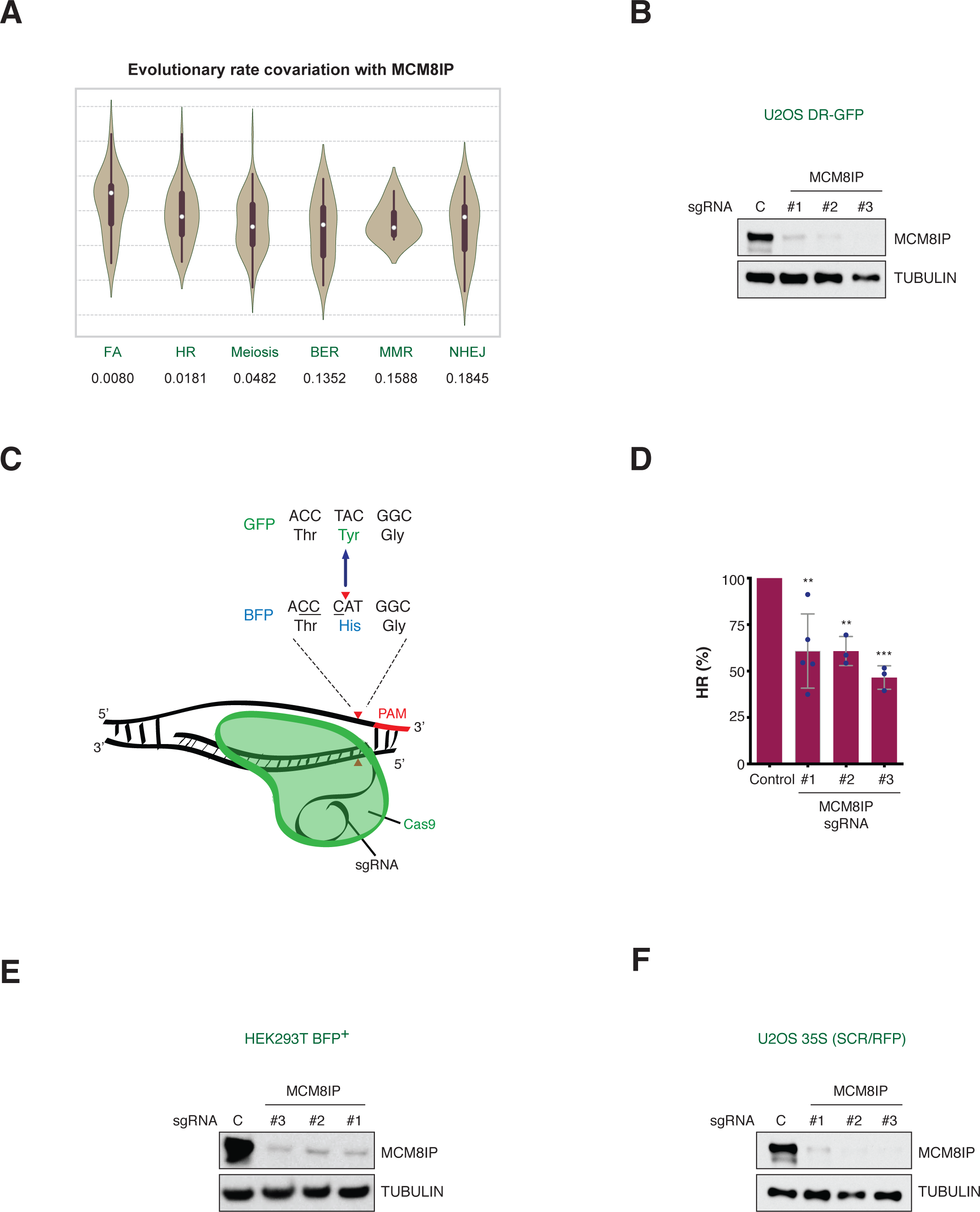
Characterization of the role of MCM8IP in homologous recombination. (A) Evolutionary Rate Covariation (ERC) analysis of MCM8IP with genes in different mammalian DNA repair pathways. Overlaid box plots indicate the quartiles of each distribution and a white dot represents the median. Permutation p-values listed below each DNA repair pathway reflect the significance of MCM8IP’s co-evolution with each pathway. FA, Fanconi anemia; HR, homologous recombination; BER, base excision repair; MMR, mismatch repair; NHEJ, non-homologous end joining. (B) Detection by western blot of MCM8IP in U2OS DR-GFP cells expressing the indicated MCM8IP sgRNAs or a non-targeting control sgRNA. Tubulin is shown as a loading control. (C) Schematic representation of the BFP gene conversion assay. Cas9 induces a DSB within the His66 codon of BFP (red arrowhead). Gene conversion results in Tyr66 and GFP expression. (D) Graphical representation of the percentage of HR events in HEK293T BFP-positive cells expressing the indicated MCM8IP sgRNAs relative to the non-targeting control sgRNA. The mean ± SD of three or more independent experiments is presented. Statistical analysis was conducted by one-way ANOVA (**p < 0.01, ***p < 0.001). (E) Detection by western blot of MCM8IP in HEK293T BFP-positive cells expressing the indicated MCM8IP sgRNAs or a non-targeting control sgRNA. Tubulin is shown as a loading control. (F) Detection by western blot of MCM8IP in U2OS 35S cells expressing the indicated MCM8IP sgRNAs or a non-targeting control sgRNA. Tubulin is shown as a loading control.

**Supplementary Figure 6.**
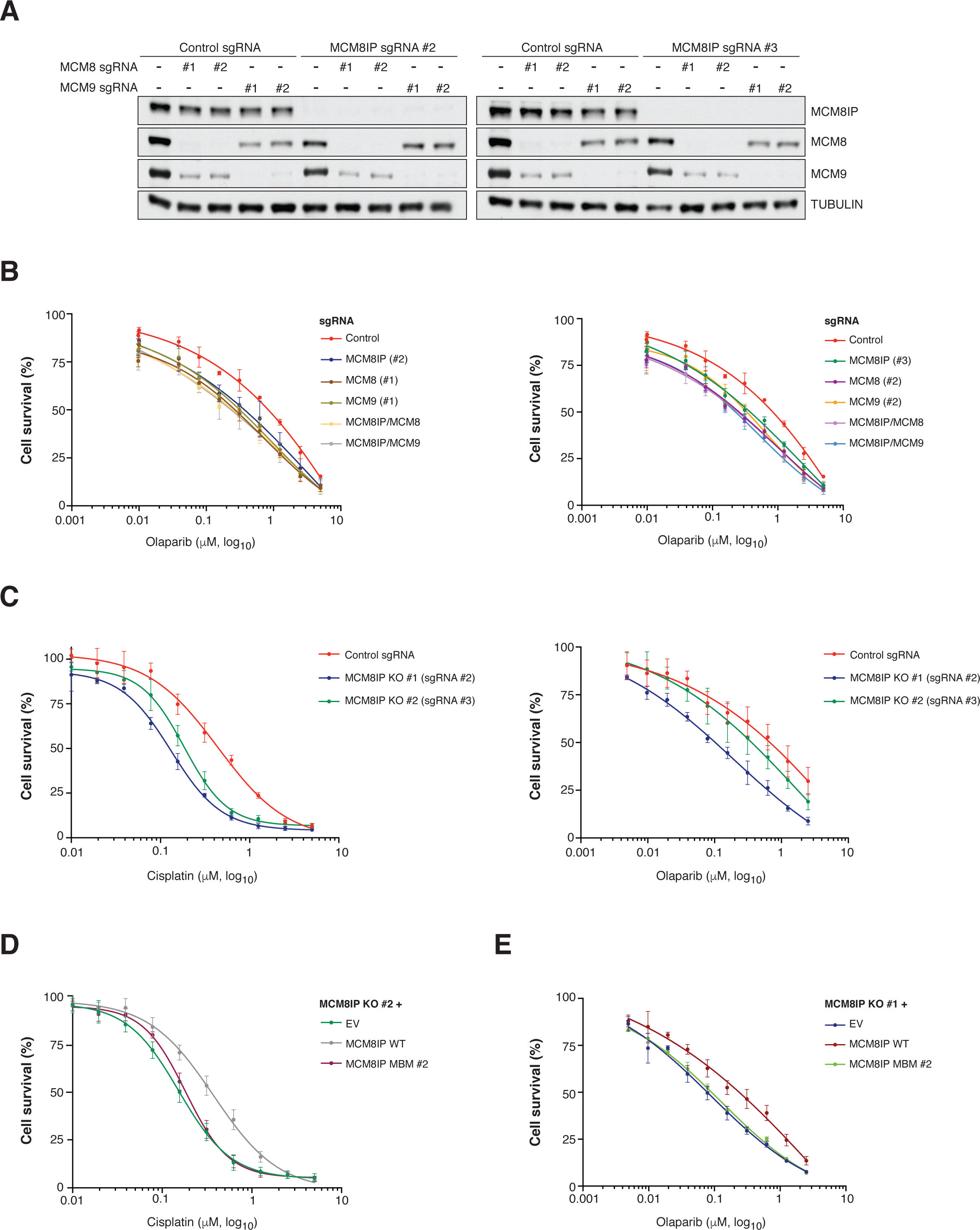
Survival analysis in MCM8IP-deficient cells upon cisplatin or olaparib treatment. (A) Detection by western blot of MCM8IP, MCM8 and MCM9 in HCT116 cells expressing the indicated sgRNAs. Tubulin is shown as a loading control. (B) Survival analysis of olaparib-treated HCT116 control cells or cells expressing either MCM8IP sgRNA #2, MCM8 sgRNA #1, MCM9 sgRNA #1 (left panel) or MCM8IP sgRNA #3, MCM8 sgRNA #2, MCM9 sgRNA #2 (right panel), alone or in combination. Cell survival is expressed as a percentage of an untreated control. The mean ± SD of three or more independent experiments is presented. (C) Survival analysis of HCT116 control cells, MCM8IP KO #1 and MCM8IP KO #2 in response to cisplatin (left panel) or olaparib (right panel). Cell survival is expressed as a percentage of an untreated control. The mean ± SD of three independent experiments is presented. (D) Survival analysis of HCT116 MCM8IP KO #2 cells reconstituted with empty vector (EV), MCM8IP WT or MBM #2 in response to cisplatin. Cell survival is expressed as a percentage of an untreated control. The mean ± SD of four independent experiments is presented. (E) Survival analysis of HCT116 MCM8IP KO #1 cells reconstituted with EV, MCM8IP WT or MBM #2 in response to olaparib. Cell survival is expressed as a percentage of an untreated control. The mean ± SD of three independent experiments is presented.

**Supplementary Figure 7.**
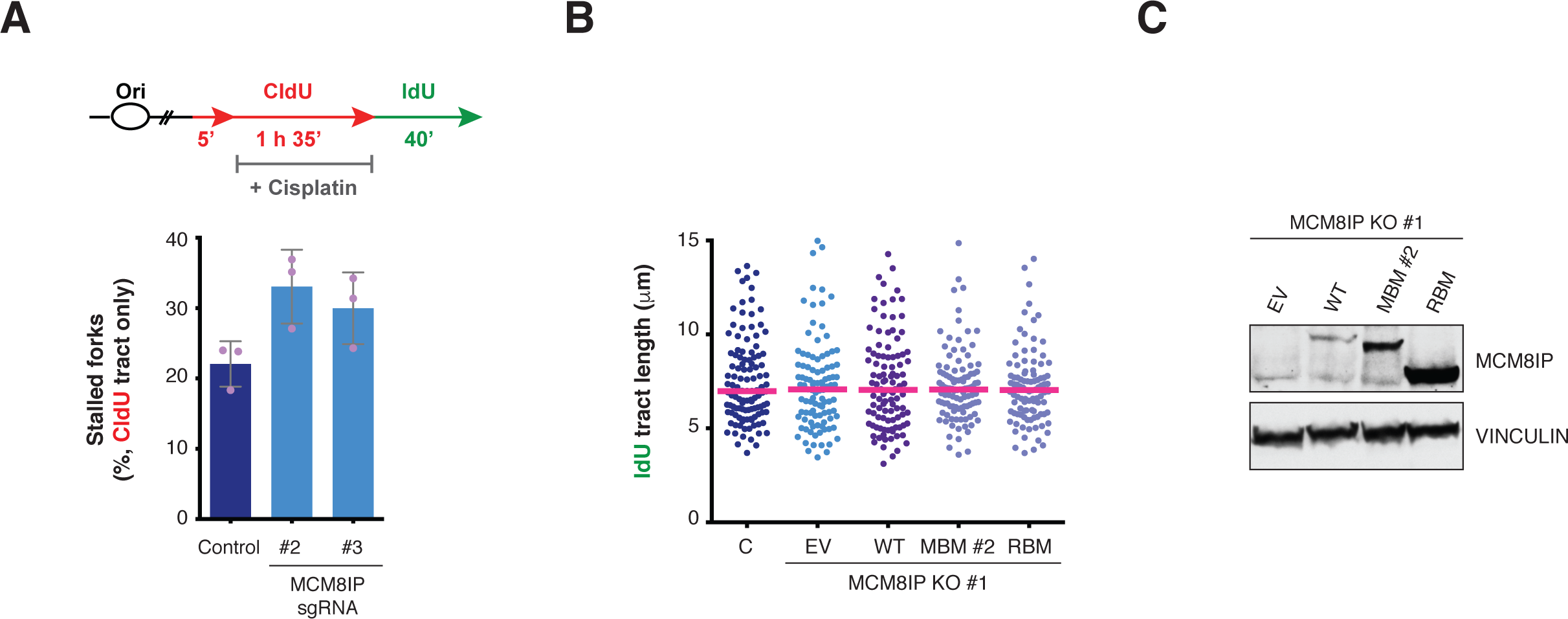
Characterization of the role of MCM8IP at replication forks in response to cisplatin treatment. (A) Schematic representation of a CldU/IdU pulse labeling assay (top panel) to assess the restart of stalled forks following treatment with cisplatin (30 µM). Graphical representation of the percentage of CldU-only tracts in cisplatin-treated HCT116 cells expressing the indicated MCM8IP sgRNAs or a non-targeting control sgRNA (bottom panel). The mean ± SD of three independent experiments is presented. (B) Dot plot of IdU tract length for individual replication forks in untreated HCT116 non-targeting control cells or MCM8IP KO #1 cells reconstituted with MCM8IP WT, MBM #2, RBM or an empty vector (EV) control. Experiments were conducted as shown in Figure 7C. Data are shown and analyzed as in Figure 7B and are representative of two independent experiments. (C) Detection by western blot of MCM8IP in HCT116 control cells or MCM8IP KO #1 cells reconstituted with MCM8IP WT, MBM #2, RBM or EV. Vinculin is shown as a loading control.

